# Product-stabilized filamentation by human glutamine synthetase allosterically tunes metabolic activity

**DOI:** 10.1101/2025.07.04.663231

**Authors:** Eric Greene, Richard Muniz, Hiroki Yamamura, Samuel E. Hoff, Priyanka Bajaj, Markus C.B. Tecson, Cyrina Geluz, D. John Lee, Erin M. Thompson, Angelika Arada, Gyun Min Lee, Massimiliano Bonomi, Justin M. Kollman, James S Fraser

## Abstract

To maintain metabolic homeostasis, enzymes must adapt to fluctuating nutrient levels through mechanisms beyond gene expression. Here, we demonstrate that human glutamine synthetase (GS) can reversibly polymerize into filaments aided by a composite binding site formed at the filament interface by the product, glutamine. Time-resolved cryo-electron microscopy (cryo-EM) confirms that glutamine binding stabilizes these filaments, which in turn exhibit reduced catalytic specificity for ammonia at physiological concentrations. This inhibition appears induced by a conformational change that remodulates the active site loop ensemble gating substrate entry. Metadynamics ensemble refinement revealed >10 Å conformational range for the active site loop and that the loop is stabilized by transient contacts. This disorder is significant, as we show that the transient contacts which stabilize this loop in a closed conformation are essential for catalysis both *in vitro* and in cells. We propose that GS filament formation constitutes a negative-feedback mechanism, directly linking product concentration to the structural and functional remodeling of the enzyme.

## Introduction

To support the metabolic needs of a cell, enzymes must be regulated to adapt to changing nutrient conditions (Atkinson 1965). Because these changes often occur faster than the rates of transcription/translation or degradation, this regulation is conferred allosterically by downstream metabolites or posttranslational modifications, or orthosterically via substrate availability and/or product inhibition (Mathy and Kortemme 2023). In metabolism, one common allosteric mechanism is feedback inhibition whereby downstream metabolites alter the enzymatic activity of upstream processes, typically at rate-limiting or committed metabolic nodes (Gerhart and Pardee 1962). In classical models of allostery, this regulation is conferred through the binding of an effector molecule at a site other than the protein’s active site, known as the allosteric site (Cui and Karplus 2008). Allosteric ligand binding subsequently induces or stabilizes a conformational change in the protein that can manifest at the tertiary, quaternary, and/or quinary (e.g. filamentous or other supramolecular states) level leading to the activity change observed (Hvorecny and Kollman 2023).

While many well-studied metabolic enzymes have evolved dedicated small molecule or protein allosteric binding sites as a primary means of regulation, an increasing number of soluble metabolic enzymes have been observed to adopt supramolecular, filamentous quinary structures as a mechanism for allosteric regulation (Noree et al. 2019; Park and Horton 2019). Filament formation can confer multiple distinctive mechanisms of enzyme regulation including repression and activation, providing an alternative to synthesis/degradation to rapidly alter the balance of enzyme activity within the cell (Lynch et al. 2020; Hvorecny and Kollman 2023). Indeed, more classic small molecule binding sites can regulate enzyme filamentation and the combination of these two modes represents a rich expansion of classical allostery, integrating structural and functional changes at the level of protein assembly.

Here, we examine how glutamine metabolism is controlled by different potential allosteric mechanisms. Glutamine levels must be critically regulated, as it uniquely serves as both an anaplerotic donor (carbon source) and a nitrogen donor in key metabolic processes (Tecson et al. 2025; Fung et al. 2025). In addition, it is, obviously, a building block in protein biosynthesis. Glutamine synthetase (GS) is an ancient enzyme that catalyzes the ATP-dependent condensation of glutamate and ammonia to glutamine (Eisenberg et al. 2000; de Carvalho Fernandes et al. 2022; Tecson et al. 2025). Glutamine is the most abundant amino acid in blood, where a majority originates from *de novo* synthesis by GS (Tecson et al. 2025; Fung et al. 2025; Castegna and Menga 2018). Moreover, ammonia generated through catabolic processes is detoxified through GS-catalyzed glutamine formation. The ammonia detoxification is critically important in the brain, where loss of function GS variants cause severe developmental defects (Spodenkiewicz et al. 2016). Regulation of human GS has so far been largely described at the transcriptional or protein degradation levels (Eisenberg et al. 2000; Rigual et al. 2025; Castegna and Menga 2018) where GS expression is restricted to certain cell types and tissues and expressed GS can be degraded by the ubiquitin proteasome system in a glutamine dependent manner (Zhao et al. 2025; Van Nguyen et al. 2016, 2017; Van Nguyen 2021). However, regulation via these processes would require hours to achieve appreciable changes in intracellular GS concentrations. By contrast, multiple allosteric and feedback mechanisms have been described for bacterial GS homologs, including by filamentation in both *E. coli* (Eisenberg et al. 2000; Huang et al. 2022) and *S. cerevisiae* (Petrovska et al. 2014; He et al. 2009). These observations led us to ask how human GS activity could be regulated on the timescales required to adapt to dynamic changes in nutrient availability.

Here, we used a combination of cryo-electron microscopy (cryoEM), enzymology, and cell proliferation assays, to discover a regulatory mechanism that incorporates both feedback inhibition and filamentation. We find that human GS forms filamentous assemblies that are stabilized by its own reaction product, glutamine. Human GS filaments display lower catalytic efficiency for ammonia, which we attribute to changes in protein flexibility surrounding the active site. We probe the importance of this change in protein dynamics with mutagenesis using enzyme activity *in vitro* and in HEK293 cells. Our findings support a model where product-stabilized GS filaments serve as an allosteric feedback mechanism, regulating activity through modulation of the enzyme’s conformational landscape.

## Results

### Identification of heterogeneous oligomeric assemblies of human GS

To define regulatory mechanisms of GS, we first used cryoEM to examine the conformational landscape of GS. Our initial attempts at *in vitro* characterization of GS using N-terminally His_6_-tagged constructs were met with low molecular weight complex compositions by size-exclusion chromatography (SEC) and we observed rare, decameric 2D averages by negative-stain electron microscopy (**Figure 1A, B**). However, most particles and 2D classes were distorted and this preparation did not result in a large population of the canonical decameric form observed in the X-ray structure of the identical GS construct (Krajewski et al. 2008). However, scarless preparations of GS, either through N-terminal His_6_-SUMO-tag and Ulp1 cleavage during purification or C-terminal Intein-CBD-His_6_ tag and DTT facilitated intein self-cleavage yielded pure enzyme with both decameric and higher molecular weight species observed by SEC (**Figure 1A, B**). In addition to scarless preparations yielding canonical forms of GS, we found a >5-fold higher *k_cat_* /*K*_M*,glutamate*_ for scarless GS, and that the His_6_-tagged counterpart has a decoupled reaction whose ADP-release was insensitive to ammonia concentration (**Figure 1C**, **Supplementary Figure 1**). These data demonstrate a significant structural and functional impact of N-terminal His_6_ tags on GS in addition to the presence of supramolecular architectures.

**Figure 1.**
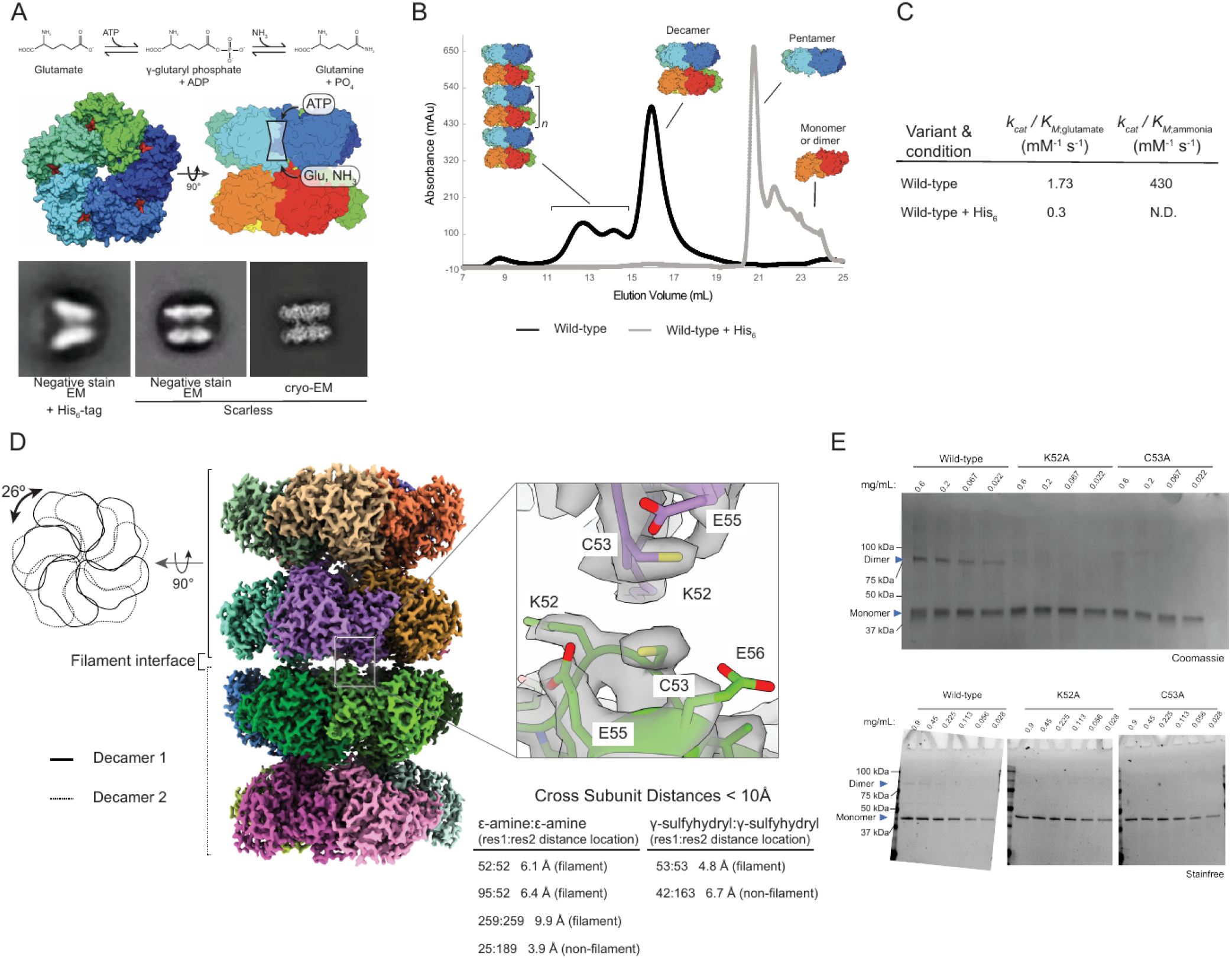
Human Glutamine Synthetase Forms Filaments. A) At top, reaction mechanism of glutamine synthetase. Middle: surface representation of PDB:2QC8 ADP + Methionine sulfoximine (MSO) bound X-ray crystal structure of human glutamine synthetase. Bottom: N-terminal His_6_-tag perturbs structure of human glutamine synthetase (GS) as observed by 2D classification of negative stain and cryoEM data. B) Size-exclusion chromatography reveals His_6_-tag dramatically alters the oligomeric state of human GS and scarless GS can form higher order oligomers. C) His_6_-tagged GS displays strongly altered steady-state kinetic constants when compared to scarless GS. N.D. = not defined as His_6_-tagged GS also display inefficient coupling of ATP hydrolysis with glutamine formation and thus *K*_M*,ammonia*_ could not be defined for this enzyme. Kinetic constants were determined from n=3 technical replicates. D) cryoEM reconstruction of apo-GS filament form. Decamers are rotated relative to each other by ∼26 degrees. The minimal five-fold filament interface formed consists of K52, C53, and E55 of adjacent monomers from different decamers. No obvious connecting cryoEM density is observed in the apo map. Cross Subunit Distances that are less than 10 between cysteine residues or lysine residues are listed as potential crosslinking sites for bifunctional crosslinkers Bis-sulfosuccinimidyl glutarate (BSG) or bis-maleimoethane (BMOE) E) Bifunctional crosslinking with BSG (top) or BMOE (bottom) to covalently attached adjacent primary amines (K52) or cysteines (C53) respectively followed by SDS-PAGE reveals crosslinked species only in the wild-type GS protein sample, but not K52A or C53A indicating presence and disruption of filament formation respectively.

To investigate how differences in oligomeric state might impact the structure and function of GS, we characterized the decamer oligomeric fraction isolated by SEC by cryoEM. We observed GS to be present in decameric form, but also a resolvable population of short filament assemblies of repeating decameric units. These higher order assemblies are consistent with dynamic exchange corresponding to the distribution of SEC peaks with estimated molecular weights of ∼0.3 - 1.5 MDa (**Figure 1B**). We isolated the 2-decameric/di-decameric particles from the decameric particles using 2D classification and non-heterogeneous refinement and reconstructed a high resolution map with FSC_0.143_ of 2.38 Å (**Supplementary Figure 2** and **Supplementary Figure 3**).

The filament was composed of two decamers (“2-decamer”) bound to each other in a ‘head-to-head’ configuration with one decamer rotated ∼26° to the next (**Figure 1D**). The decamer-decamer (filament) interface is five-fold symmetric and made up of residues K52, C53, and E55 contributed by both monomeric subunits across the interface. Interestingly, K52 also participates in a binding interface between GS and bestrophilin2 ion channel (Owji et al. 2022). We observed little to no density at the interface despite the stable relative orientation of both decamers to each other (**Figure 1D**). These data suggest that the formation of the interface is concentration dependent and driven primarily by electrostatic interactions. Alternatively, the interface could be largely solvent-mediated or be connected by a ligand(s) that occupy only some of the 5 symmetry related interfaces. In both scenarios, any signal is likely to be averaged out and experiments with symmetry expansion and focused classification did not yield any convincing density.

To test the importance of the interface, we used homo-bifunctional crosslinking to probe the contributions of individual contacts. The 2-decameric map suggests that only one pair of lysine residues (K52:K52) and one pair of cysteine residues (C53:C53) are surface exposed and within <10 across two monomers. To ensure a stringent selection for only K52 and C53 we used either the 7.7 spacer-arm bifunctional NHS-ester crosslinker (bis(sulfosuccinimidyl) glutarate or BSG) or the 8.0 spacer-arm biofunctional maleimide crosslinker (bismaleimidoethane or BMOE). Crosslinking of two monomers by either crosslinker was observed in the wild-type enzyme and more prevalent at higher enzyme concentrations (**Figure 1E**). However, mutation of either K52 or C53 to alanine ablated the crosslinking reactivity and reduced filament formation as assessed by negative stain EM (**Supplementary Figure 5**). Together these data support the 2-decameric filament interface assignment whose formation is concentration dependent and driven primarily by electrostatic interactions.

### Glutamine stabilizes larger GS filaments

The interface defined by the 2-decameric assembly and the presence of higher molecular weight species in the SEC trace suggested that larger filaments of GS can readily form. Next, we wanted to understand what functional consequence may be conferred by filamentation in human GS. We asked if GS filament formation could be retained under turnover conditions and impact the catalytic function of GS. CryoEM is an ideal tool to investigate structural changes to macromolecules under turnover conditions because substrates can be added to the enzyme immediately before vitrification freeze-quenching to preserve the sample in a near-native context (**Figure 2A**, **Supplementary Figure 6**). Turnover conditions were induced by adding saturating concentrations of substrates immediately prior to vitrification, with reaction times totalling between 49 and 59s initiated by manual mixing prior to blotting. Our initial grid screening of GS under turnover conditions identified filaments at lower GS concentrations (0.1 mg/mL) than we observed for apo GS. Initial reconstruction of these screening data revealed two new structural features in the GS filament: defined EM density between both decamers and missing density for the active site loop region (**Figure 2A**).

**Figure 2:**
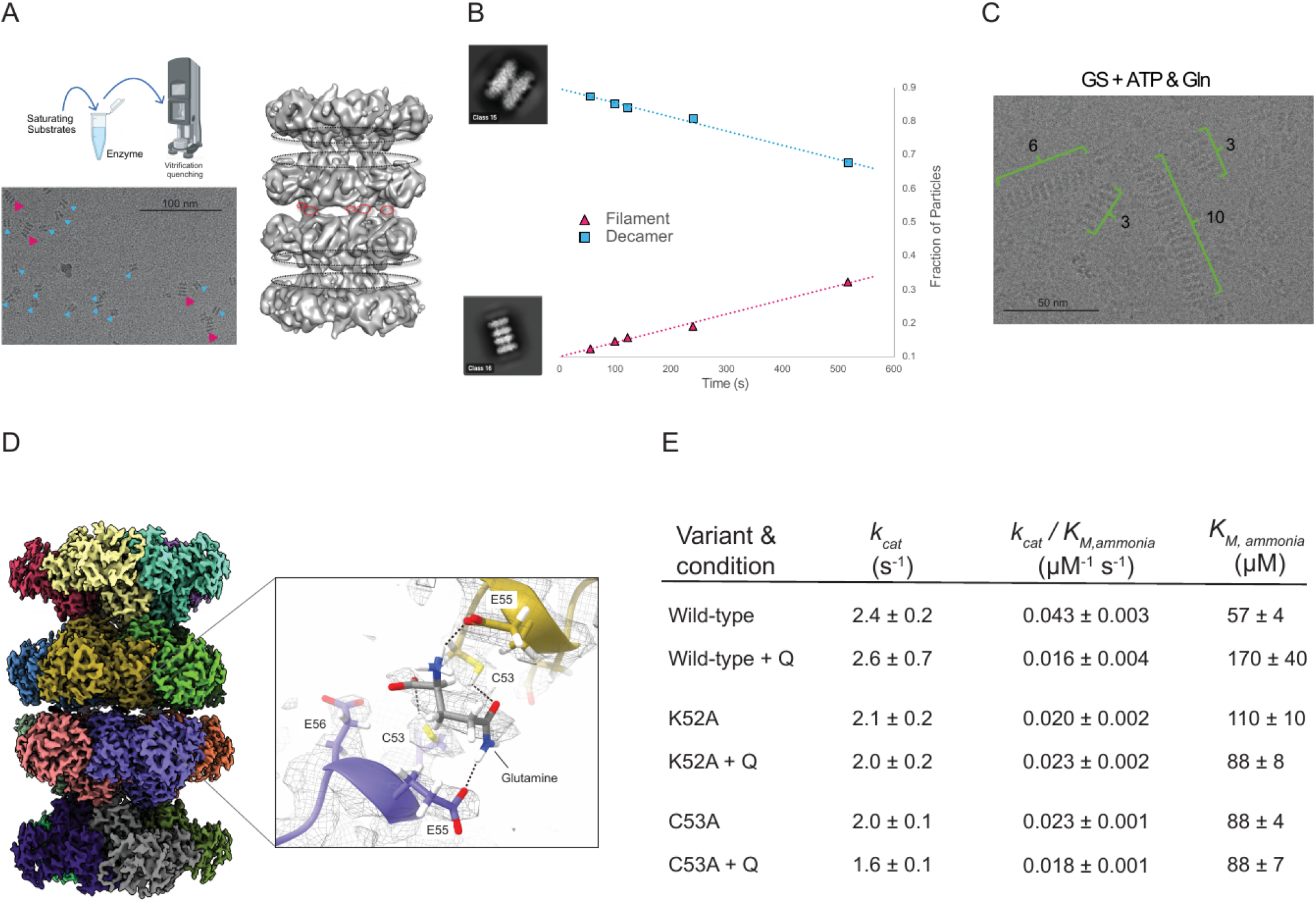
Glutamine binds to GS filaments at the interface. A) Establishment of steady state conditions prior to vitrification with GS at low concentration (0.15 mg/mL) revealed presence of 2-decameric species (red arrows) in addition to decameric species (blue arrows) during cryoEM grid screening. Initial reconstruction of the turnover 2-decameric map demonstrated clear density connecting decamers (red dotted circle) not previously observed in the apo-filament map and loss of E305-loop density (black dotted circle). B) Time-resolved cryoEM demonstrates linear relationship between glutamine and filament formation assessed by independently processing five datasets and quantifying the fraction of decamer and filament present (see Methods) and plotted against the vitrification quench time of the reaction. C) cryoEM screening data demonstrating robust filament seeding from addition of ATP and glutamine to human GS prior to vitrification. D) Turnover-filament global 3D reconstruction (left) and locally refined filament interface (right) demonstrates similar overall architecture to apo-filament map (left) but with clear density between decamers with glutamine modeled which forms hydrogen bonds with K52, C53, and E55 (right). E) Steady-state ammonia kinetic constants for wild-type, K52A, and C53A with and without 10 mM glutamine added to reaction to stimulate filament formation. Errors were calculated as S.E.M. from global fitting and were propagated through each Michaelis-Menten parameter (Equations 2-4; *n* = 8 technical replicates) see Methods.

Given that the only sample preparation differences between the apo-GS datasets and the turnover conditions was the addition of substrates that are converted to products prior to vitrification, we hypothesized that one or multiple substrates (ATP, glutamate, and ammonium) or products (glutamine, PO_4_, and ADP) may be responsible for this additional density. However, at this initial low resolution we could not assign a compound at this interface nor understand if an interface-bound compound had any conformational impact on active site geometry.

To discriminate whether the additional filament density corresponded to product or substrate, we performed a time-resolved (tr-)cryoEM experiment. We collected five turnover cryoEM datasets that differed only in the vitrification-quench time of the steady-state reaction with reaction times between 57s and 517s corresponding to high substrate and high product respectively (see methods for details) at a fixed concentration of enzyme (0.3 mg/mL). To understand if there was a trend in the particle populations between the decamer/filament compositions as a function of reaction time, we compared the number of particles in decameric versus filamentous states by 2D classification (**Supplementary Figure 7**). The same template picking parameters were applied for each tr-cryoEM dataset and 2D classification was performed thrice; once with each time-point dataset processed independently and twice where all particles were combined and the dataset identity only revealed after 2D classification particle classes were picked and 2D classes were picked with and without top views (**Supplementary Figure 9** and **Supplementary Figure 10**). We then normalized decamer and filament particle picks to the total number of picked particles (excluding 2D classes that could not be assigned either a decamer or filament) and plotted these as a function of reaction time (**Figure 2B**). The positive linear correlation between fraction of particles in a filament-like assembly and reaction time suggests that a product of the forward reaction of GS acts to stabilize the filament assembly. Subsequent screening with high concentrations of glutamine revealed large >4 decameric unit filament assemblies (**Figure 2C**), suggesting that product formation has a positive effect on filament assembly.

To define how products stabilized the GS filament assembly, we next determined a high resolution structure of the filaments imaged under turnover conditions. We performed an additional cryoEM experiment with increased GS concentration (1.5 mg/mL), to bias our particle populations towards filamentous assemblies, and vitrified at 57s, a reaction time that should yield a high concentration of product due to the higher enzyme concentration than previous experiments. The resulting 3D reconstruction of the 2-decameric species had an FSC_0.143_ of 2.2 (**Supplementary Figure 11**A and **Supplementary Figure 8**).

Inspection of the ‘filament assembly’ interface between the two decamers revealed a discrete density that was absent from the apo-filament reconstruction (**Figure 2D**). The density is too small for a nucleotide (ATP or ADP) and too large for inorganic phosphate (**Supplementary Figure 11**B). Given the association with product formation, we modeled glutamine into this density in a conformation consistent with forming hydrogen bonds that link the filament interface residues and the bound product (**Supplementary Table 2**). The two-fold symmetry at this interface and the averaging inherent in our data processing results in density that likely reflects contributions from glutamine in two orientations. To relieve symmetry averaged artefacts, we symmetry expanded particles and subjected them to focused refinement on a single binding site with a mask covering four subunits total to support particle alignment (**Figure 2E**). The resulting density was supported by glutamine positioning similar to the symmetry refined global map (**Supplementary Table 2**). In addition, we note two potential, and not mutually exclusive, reasons we were able to observe filaments in the apo enzyme. First, we used high concentrations of GS, which likely shifts the equilibrium towards the filament form even in the absence of glutamine. Second, there may be a small, unresolved, amount of glutamine that persists during purification that occupies only a minor fraction of interfaces per di-decamer. As discussed above, such contributions would be averaged out during map refinement in the scheme we have used here. Taken together, these data support a model wherein glutamine synthetase filamentation generates a composite binding site for glutamine at the filament interface, which is located over 10Å from the active site.

### Glutamine stabilized filaments have lower catalytic efficiency for ammonia

We next examined the functional consequences of GS filamentation on enzymatic activity. To assess steady-state kinetic differences between decameric and filamentous compositions of GS, we assayed different isotropic peak fractions resolved by SEC immediately after separation. We found *k_cat_* and *k_cat_* /*K*_M_ for all substrates were completely in agreement between the decamer and 2-decamer fractions at 0.01 mg/mL for wild-type, K52A, and C53A (**Supplementary Figure 13**). However, this enzyme concentration is 30-fold dilute compared to typical cryoEM conditions and the 2-decameric fraction significantly decomposed into decamer, as assessed by mass photometry (**Supplementary Figure 14**A). Upon increasing enzyme concentration to 0.1 mg/mL, which is within the range of our cryoEM experiments (0.3 mg/mL for all datasets except where indicated) and shown to support filamentation by chemical crosslinking (**Figure 1D**), we uncovered a subtle, 2-fold *K*_M,_ *_ammonia_* defect for the 2-decamer fraction compared to the decamer fraction for wild-type GS (**Supplementary Figure 14**B).

To probe this defect in K_M_ further, we took the decameric fraction at 0.1 mg/mL and repeated steady-state assays in the presence or absence of the glutamine concentration used to stabilize filamentation by CryoEM (**Supplementary Figure 15**). The presence of glutamine increased *K*_M,_ *_ammonia_* three-fold for WT, but had smaller effects for the mutations at the interface that directly interact with the ligand (**Figure 2F**). Given the incomplete filament formation observed in our cryoEM experiments, with a maximum filament formation of ∼50% at 0.3 mg/mL concentration (**Supplementary Figure 9**), the effective *K*_M,_ *_ammonia_*defect upon filamentation is likely at least 2-fold higher than measured. The range of *K*_M,_ *_ammonia_* values reported here are near the ammonia concentration estimates reported in the blood and liver under normal physiological conditions (<50 μM) and lower than hyperammonemic conditions (0.2 - 1 mM) (Albe et al. 1990; Dasarathy et al. 2017).

### The filament state impacts conformational variability of E305-loop

To understand the structural basis underlying the *K*_M,_ *_ammonia_* defect, we inspected the interactions of active site residues and loops for differences across the oligomeric forms of GS. The GS active site forms a bifunnel shape with ATP gaining access to one subsite with glutamate and ammonia accessing the other subsite (**Figure 3A**). The glutamate/ammonia binding site is gated by a flexible loop that bears two key catalytic residues: R299, which interacts with glutamate substrate carboxyl group, and E305, which is predicted to deprotonate ammonium ions to ammonia, enabling the required nucleophilic attack of the y-glutamyl phosphate intermediate (Owji et al. 2022; Krajewski et al. 2008). Previously, crystal structures of human GS revealed a closed loop conformation in the presence of the transition state inhibitor methionine sulfoximine (PDB 2QC8). However, this loop exhibited weak electron density in the ADP-bound state (PDB 2OJW), which potentially indicates mobility of this element throughout the catalytic cycle.

**Figure 3:**
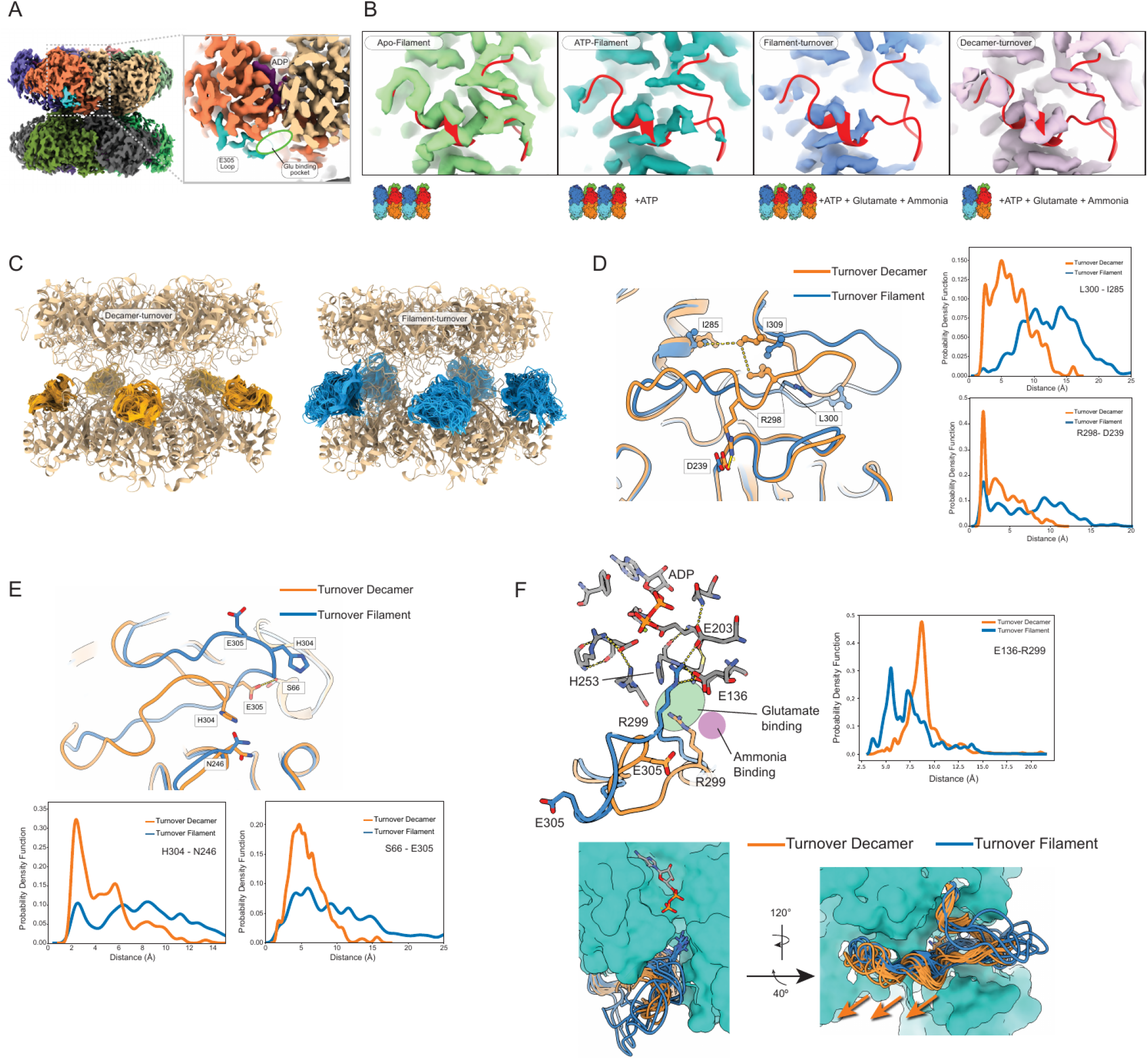
Variable E305-loop density and conformations in human glutamine synthetase. A) Turnover-decamer cryoEM density map is colored based on monomer except for one E-305 loop with is colored cyan and ADP density that is colored purple demonstrating how the E305-loop closes over the active site to enclose the glutamate and ammonia binding site. B) CryoEM E305-loop density representations for the apo-filament, ATP-filament, turnover-filament, and turnover-decamer maps with 2QC8 docked into each map shown in red. The contour is chosen to equalize the relative density levels in other regions of the protein (**Supplementary Figure 16**). C) Representation of EMMIVox ensembles of the E305-loop (residues 295-309) for one pentamer (decamer-orange and filament-blue) overlaid with ribbon structure of consensus model (tan). D) Single filament and decamer turnover monomers overlaid and presented in blue and orange respectively presenting a hydrophobic stabilizing patch (I285, L300, and I309) and stabilizing salt-bridge (D239 and R298) in the ‘closed active’ loop state that is missing in the filament models. Distance measurements calculated across full EMMIVox ensembles represented as kernel density function between residues participating in ‘closed active’ loop state are presented demonstrating bias away from these contacts in the turnover filament form. E) Same as (D) but representing the ‘tip’ of the loop showing E305 hydrogen bonding with S66 and H304 hydrogen bonding with N246 with quantification of these distances from the ensemble. In all cases, the filament oligomeric state under turnover conditions disfavored these interactions. F) MD-refinement of the cryoEM filament turnover map revealed active site loop infiltration into the active site with R299 hydrogen bonding with active site residues E136 (top left) with distance quantification of R299 to E136 in the active site (top right). At bottom, overlay of all ten monomers per MD-refined model with loops in ribbon, ADP in stick, and the rest of the model is represented in surface. The filament loop is in an outward conformation from the active site at the base of helix 8.

Given the importance of this loop in gating glutamate and ammonia access to the active site, we examined how the E305-loop conformation and dynamics varied upon filament formation and substrate identity. All maps had similar resolution range (FSC_0.143_ between 1.95 - 2.27Å, **Supplementary Figure 3**, **Supplementary Figure 4**, **Supplementary Figure 8**, **Supplementary Figure 12**) and showed defined secondary structure and amino acid side chains in helix-12 (266-288) and beta sheet-11 (313-316) on either side of the loop at similar density contours (**Supplementary Figure 16**). The apo-filament cryoEM map showed well resolved density for the E305-loop with clear side chains, which support building an atomic model in a conformation we call the “closed-active state” (**Figure 3B**). Loop conformational heterogeneity was also influenced by nucleotide state, as the filament map with ATP only displayed partial density (**Figure 3B**). By contrast, the filament map collected during turnover displayed almost no density in the region, which is indicative of a greater degree of disorder. However, the E305-loop was partially restored in the cryoEM map of the turnover-decamer map (**Figure 3B**). The differential loop density between turnover-decamer and turnover-filament species was further supported by 3D classification of symmetry-expanded particles, which recovered partial E305-loop density in 4/12 turnover-decamer classes (C5; **Supplementary Figure 18**) compared to 0/12 classes for the turnover-filament (D5; **Supplementary Figure 19**). This observation implicates the E305-loop conformational distribution as a potential factor contributing to the observed impaired kinetics of GS filaments. Taken together the E305-loop is sensitive to nucleotide state and also by filament state under turnover conditions.

The changes in density for the loop results from conformational heterogeneity that is ‘averaged out’ upon reconstruction. To provide an atomistic model of the conformational space explored by the E305-loop, we used Bayesian inference to balance experimental cryoEM data with accurate physico-chemical molecular dynamics as implemented in EMMIVox ensemble refinement (Hoff et al. 2024). Across different conditions, the conformation of the loop was dominated by the closed-active loop conformation (**Figure 3C**). However, the turnover-filament model displays distended loop conformations diverged from the active-closed state that we turn collectively as the ‘open ensemble’ of loop conformations (**Figure 3C**). The E305-loop (residues 299 - 311) sampled a more diverse ensemble (with root mean square fluctuation values of greater than 10Å) in the turnover filament state, with the turnover decamer and apo-filament displaying reduced variability (**Supplementary Figure 17**). By contrast, across all states, the remainder of the active site maintained similar fluctuations (**Supplementary Figure 17**).

We next examined the principal contacts that serve to stabilize the ‘active closed’ state of the loop (**Figure 3D,E**). We observe multiple residue contacts that appear stabilizing including a hydrophobic pocket formed by L300, I285, and I309, a salt-bridge formed between R298 and D239, and a hydrogen bond between H304 and N249 (**Figure 3D,E**). All these interactions displayed favored distances in the decamer-turnover compared to filament turnover ensembles. A major consequence of this increased loop mobility is that Arg299 can “invade” the active site (**Figure 3F**). We quantified the distance distribution between R299 and E136 in our turnover-decamer and turnover-filament ensembles and found a substantial bias towards R299 infiltration of the active site in the filament ensemble that is absent in the turnover decamer ensemble (**Figure 3F**). The conformation of Arg299 could potentially sterically hinder substrate access to the active site and thus contribute synergistically with loop mobility to the observed *K_M, ammonia_* defect. Collectively, these results suggest that biasing of conformational distribution of the loop is important for tuning activity in GS.

### Global conformational change coordinates E305-loop dynamics

We next asked what conformations or structural features underlie these changes to E305-loop conformational space and how filament formation promotes an ‘open ensemble’. Utilizing our EMMIVox refined models as a starting point, we assessed global conformational changes between the turnover-filament and turnover-decamer structures that might influence the local conformation of the loop. First, we found a rotational conformational change (1.2°) apparent about the pentamer-pentamer interface (**Figure 4A**). Given that this conformational change could impact the local conformation at each composite active site by altering monomeric interfaces we explored how this global conformational change related to the overall conformation of each monomer. We observed a ‘crimping’ conformational change per monomer where in the filament structure the β-grasp domain and the catalytic domain are rotated towards each other and away from their respective active sites (**Figure 4A**).

**Figure 4.**
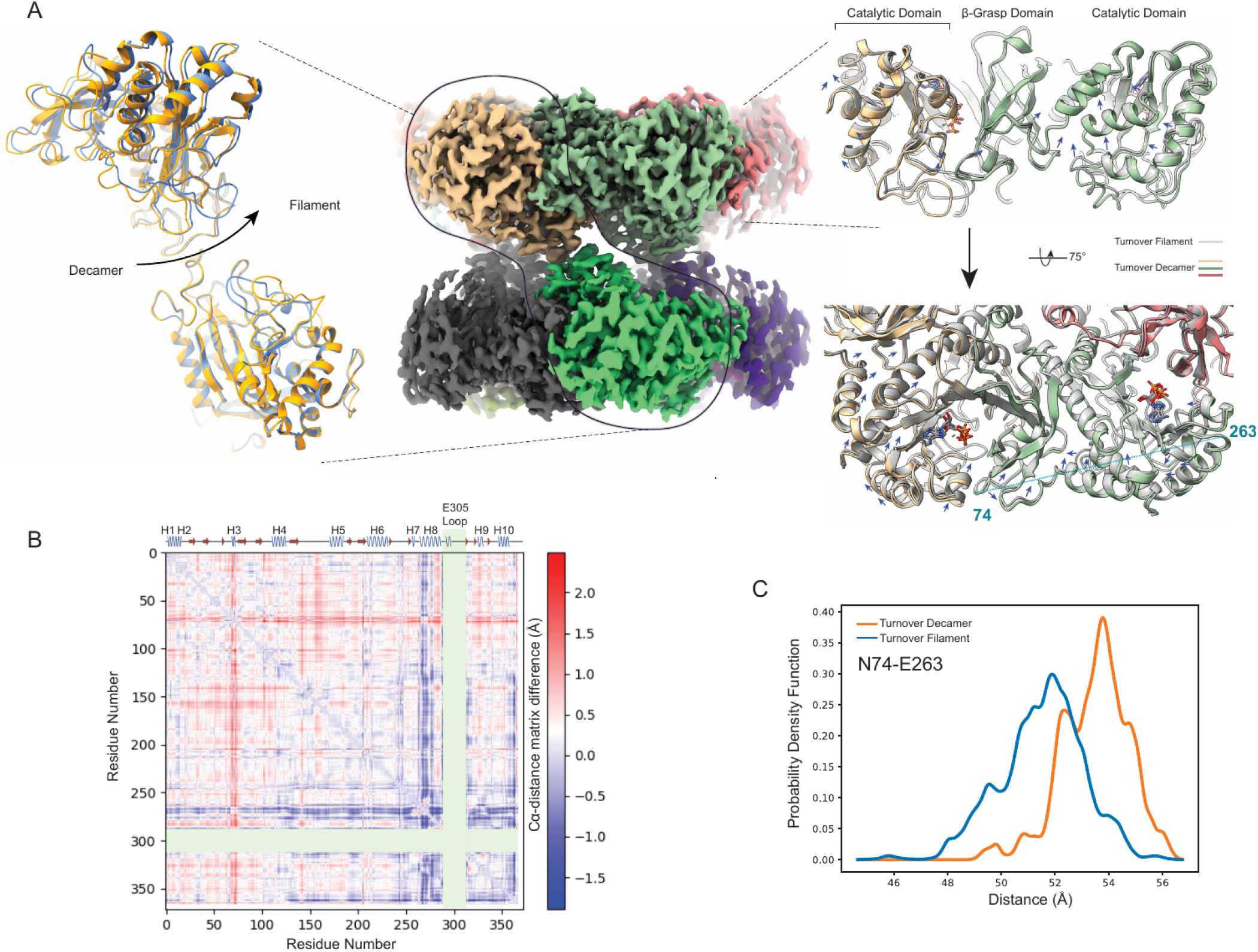
Global conformational change is associated with loop conformational heterogeneity under turnover conditions. A) Left: rotational motion about the pentameric interface observed where models were aligned on the bottom monomer and rotation seen in the upper monomer is noted (decamer in yellow and filament in blue). Middle: representation of the decameric turnover cryoEM map colored per monomer with outlines showing relevant areas shown at left and right. Right: monomeric interface within a pentamer overlay of the turnover filament model (gray) with turnover decamer model colored by monomer (tan, green, and red). Ligands are colored by atom type. Arrows indicate conformational change from filament to decamer state. The largest Ca distance change not including the E305-loop is indicated between residues 74 and 263 from across the pentamer. B) Ca distance difference matrix for chain G between turnover decamer and turnover filament models with the E305-loop (residues 289-312) and the far C-terminal peptide (369-373) omitted with a cartoon secondary structure representation of a monomer on top and distance color legend at right demonstrating that the tip of the B-grasp region (60-76) and helix 8 (residues 262-282) showing the largest conformational differences. C) Ca distance differences for residues 74 and 263 between the turnover filament and turnover decamer ensembles demonstrating a robust 2 angstrom distance difference.

We next sought to define the regions within each monomer most altered upon filament formation by performing a Cα-distance matrix subtraction analysis (**Figure 4B**; **Supplementary Figure 20**). As expected, the E305-loop conformational heterogeneity dominated this analysis, however, we also observed an ∼2Å shift in the tip of the β-grasp domain (residues 70-76) and helix-8 which abuts the active site and immediately precedes the E305-loop (**Figure 4B**; **Supplementary Figure 20**). We found the largest Cα-distance change between these two regions, specifically, between N74 and E263 displaying a distance change of 2.05Å (**Figure 4A**). Finally, to understand how prevalent this global conformational change is we quantified this Cα-distance in the full turnover-filament and turnover-decamer ensembles and found persistent distance distribution changes of ∼2Å between these two ensembles (**Figure 4C**). Thus, we propose that this global conformational change serves to influence E305-loop conformational distribution via two primary mechanisms via helix 8 repositioning away from the active site in the filament form which: 1) hinders E305-loop ability to completely close over the ammonia binding site (residues S66, S67, and D76;(**Figure 4B,C**) and 2) disfavors E305-loop contacting closed-state stabilizing residues (**Figure 3D,E**).

### E305-loop conformational space is crucial for Michaelis-complex formation

Since all ensembles, including the apo-filament, could sample both the ‘active closed’ and ‘open-ensemble’ states, we next pursued mutagenesis to more directly test the importance of the loop flexibility on catalysis. We created alanine mutants of residues D239, R298, L300, I309, H304, and E305. Each of these residues reside on the E305-loop with the exception of D239 which participates in a salt-bridge interaction with R298 on the loop. E305 was chosen to be an active site control, which does not appear to stabilize the closed state of the loop, but rather, the closed state of the loop positions E305 near the proposed ammonia binding site (Moreira et al. 2017; Issoglio et al. 2016; Frieg et al. 2016).

We measured the steady-state Michaelis-Menten parameters of all variants (**Figure 5A**). The E305A variant principally impacted *K*_M,_ *_ammonia_* (∼10 fold) as would be expected from this residue’s predicted role of deprotonating ammonium ions in preparation for nucleophilic attack. The E305A variant was also the only variant to display a large *k_cat_* effect (∼10 fold) which is also expected given its presumed role in priming ammonium to ammonia in the active site (Moreira et al. 2017; Issoglio et al. 2016; Frieg et al. 2016). The *K*_M,_ *_ammonia_*parameter was also impacted for all other loop variants tested by 2-10 fold. However, the principal kinetic impact for other loop variants was on the *K*_M,_ *_glutamate_*. The magnitude of *K*_M,_ *_glutamate_* change ranged from 100 fold in L300A to 5 fold in H304A. Disruption of the hydrophobic interface of L300, I309, and I285 appeared to have largest impact on *K*_M,_ *_glutamate_*, followed by disruption of the R298*D239 salt bridge (50-fold change for both variants), followed lastly by disruption of the H304*N249 hydrogen bond.

**Figure 5.**
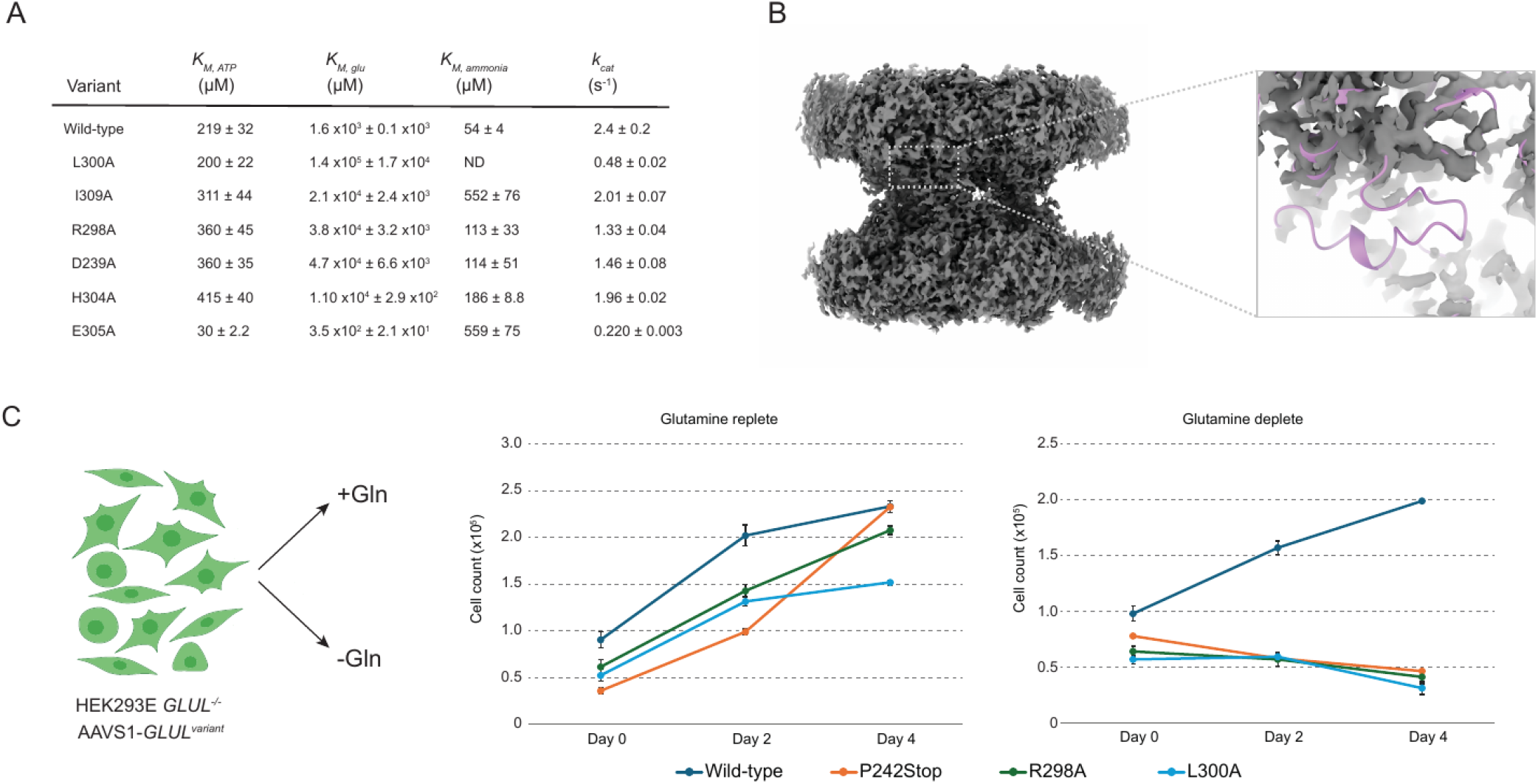
Glutamine synthetase E305-loop conformation controls Michaelis-complex formation. A) Steady-state kinetic parameters for wild-type GS (also displayed in Figure 2) and six point-mutants of GS, L300A, I309A, R298A, D239A, H304A, E305A as assessed by the coupled-ADP release assay (methods). Kinetic constants were determined from n=3 technical replicates. (B) cryoEM density map for R298A decamer under turnover conditions displayed in full (left) with inset focused on loop density (right) demonstrating complete loss of the E305-loop for this variant. (C) Schematic of *HEK293E GLUL-/-* cell line auxotrophic for glutamine is reconstituted with single genomic integration of individual *GLUL* variants (methods) and used to assess glutamine prototrophy. All variant cell lines grow equivalently in glutamine replete media conditions (middle) but only the wild-type *GLUL* variant could support cell growth and survival under glutamine deplete conditions, whereas R298A, L300A, and a premature stop codon at P242 all led to cell death. Cell proliferation assays were performed in technical quadruplicate; a single clone of the landing pad cell line was generated, and a single population of each GLUL variant cell line was created for testing.

The *K*_M,_ *_ammonia_* was impacted but generally less so than *K*_M,_ *_glutamate_* in E305-loop variants. This result likely due to the use of saturating glutamate concentrations required to estimate *K*_M,_ *_ammonia_* where the majority of active sites are glutamate bound, or y-glutamyl phosphate formed, which will have had already promoted the closed-active loop state via amino acid carboxylate-R299 interaction, thus restricting the potential scope of loop conformational variability. Even with such a presumed stabilization, the full Michaelis-complex formation is still impacted for the final ammonia substrate binding which is likely a consequence of the biased conformational state of the E305-loop. Taken together, these data indicate that mutations outside the active site to the E305-loop stabilizing interactions have large impacts to the *K*_M_ for substrates that bind to half-active sites gated by the loop. These mutations do not have pronounced effects to the overall *k_cat_* however, indicating that while Michaelis-complex formation is hindered, the rate-limiting step of catalysis is unimpacted.

To test the model that loop biasing impacts *K*_M_ values for GS, we structurally characterized the decameric form of loop variant R298A under turnover conditions. The final reconstruction yielded an overall map of FSC_0.143_ of 1.95 Å with high local resolution values across the map (**Supplementary Figure 21**). Importantly, this map, despite being the highest global resolution map reconstructed, did not display any appreciable E305-loop density (**Figure 5**), similar to the turnover filament (**Figure 3B**). Thus, the R298A mutant, even in the decameric form, would rarely adopt the active-closed conformation, explaining the *K*_M,_ *_glutamate_* and *K*_M,_ *_ammonia_* defects resulting from this mutation. Collectively, these data suggest a connection between loop ensemble and catalytic activity for human GS that is principally manifested in *K*_M,_changes for amino acid and ammonium substrates.

### Loop variants are unable to support glutamine auxotrophy

Since our data suggested that altering loop dynamics via mutation or filamentation could alter *K*_M_, we next wanted to test the effect of perturbing the loop on the overall cellular activity of GS. Most human cells require exogenous glutamine to support growth in tissue culture unless they express GS. Indeed, GS expression has been used as an auxotrophic marker for antibody production HEK293E GLUL-/- cell line (Yu et al. 2018). Starting from this auxotrophic HEK293E GLUL -/- cell line, we used CRISPR to introduce an attP landing pad (Matreyek et al. 2017, 2020), to facilitate stable cell line generation expressing individual GS variants. To create each mutant line, we transfected with Bxb1 and cognate attB GLUL variant DNA. GS variants were then overexpressed from a pTet promoter driven by doxycycline and tested for growth in the presence and absence of glutamine supplementation (**Figure 5C**)

Glutamine prototrophy of this cell line was confirmed as high cell proliferation was observed for wild-type GS in glutamine deplete media while there was a lack of survival for a variant bearing a premature stop codon inserted at P242 (**Figure 5C**). We assessed glutamine auxotrophy for GS filament interface variants C53A and K52A (**Supplementary Figure 23**). As expected, in the HEK293 model, both variants supported glutamine auxotrophy to similar levels as wild-type. This system is not likely to be sensitive to the feedback inhibition mediated by filamentation to reduce glutamine production as ammonia levels in HEK293 cells typically exceed those levels seen in healthy tissues by up to 100-fold (Rajendra et al. 2011). However, the two loop mutants, R298A and L300A, could not support growth under glutamine-deplete conditions displaying the same phenotype as P242stop. The *K*_M,_ *_glutamate_* for R298A and L300A are outside the normal steady state concentrations of intracellular glutamate, suggesting that loop conformational control of the Michaelis-complex can occur in the cell to directly impact metabolic flux through GS.

## Discussion

We propose a model for glutamine synthetase regulation through filament formation that alters the conformational landscape of the active site gating E305-loop (**Figure 6**). We show that filament formation is driven both by higher enzyme concentrations and glutamine binding at the filament interface. While the enzyme concentrations to achieve robust filamentation in the absence of glutamine are much higher than observed in cells, the protein concentrations used in our time-resolved cryoEM experiments where filamentation is correlated with accumulation of glutamine are within the range of intracellular GS concentrations in S. cerevisiae (Engel et al. 2025) and human cell lines (Wiśniewski et al. 2014). Additionally, we found that physiological concentrations of glutamine can stabilize filamentous GS structures.

**Figure 6:**
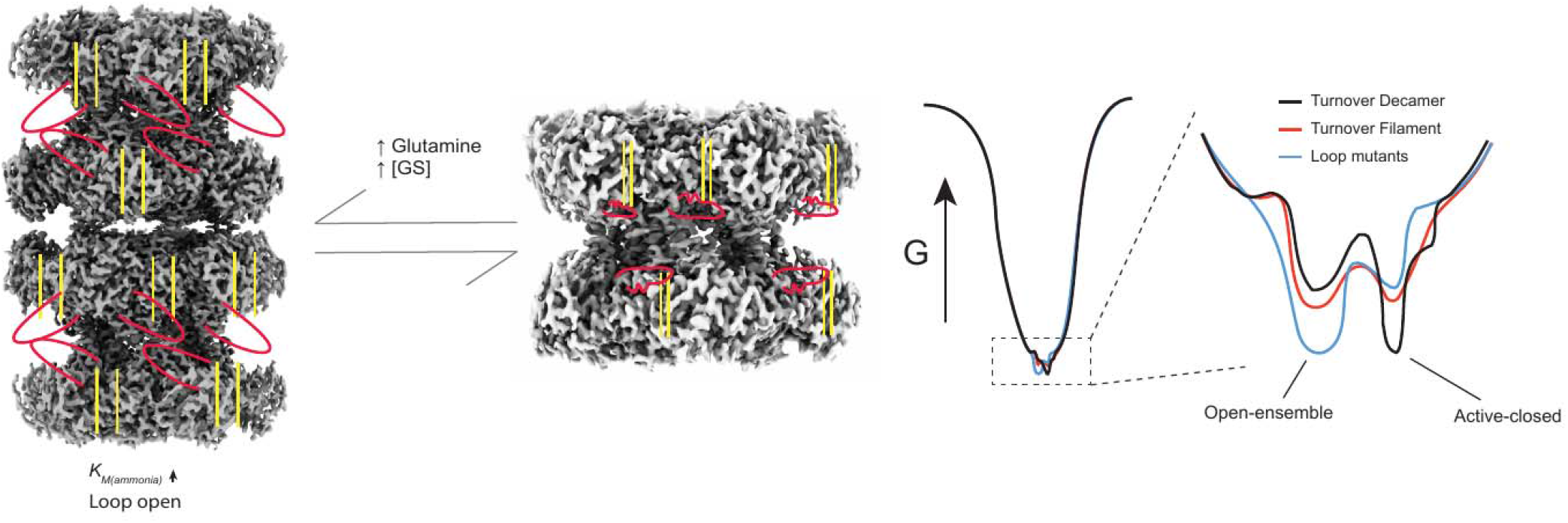
Filament formation tunes E305-loop dynamics to modulate enzyme activity. Left: cryoEM map of the turnover filament which includes a higher *K*_M,_ *_ammonia_* and open E305-loop indicated at bottom and by an ‘open’ red colored E305-loop cartoon. Also indicated is the global conformational change that opens up the monomeric interfaces about the active site (yellow lines). This form of the enzyme is increased with increased GS concentration and/or glutamine concentration and is different from the decamer which includes a more closed active site and less conformationally heterogeneous E305-loop. Right: proposed conformational landscape of GS wherein the global fold of the enzyme is largely retained and subtle conformational/ensemble changes found in the low thermodynamic basin are present. Inset: the E305-loop ensemble is depicted for the turnover decamer, turnover filament, and E305-loop mutants wherein the decamer slightly favors an active-closed state, the filament an open-ensemble, and loop variants significantly favor the open-ensemble state.

GS is an ancient enzyme that is found in all domains of life as it serves a critical crossroads between carbon and nitrogen metabolism. Given its centrality to numerous metabolic pathways, GS is subject to sophisticated intracellular regulation across the domains of life. E305-loop flexibility has been noted in X-ray crystallographic structures of human GS (Krajewski 2008) and in other homologs too (Tecson et al. 2025) supporting a model of functional flexibility of this structural element. The most well studied is type I-beta GS (GSIB) found in gram negative bacteria which contains a ‘regulatory loop’ that abuts the E305-loop at the active site. The regulatory loop can be covalently adenylated by a bifunctional enzyme that is sensitive to glutamine levels and adenylation inhibits GS (Eisenberg et al. 2000). Given our demonstration of the importance of maintaining optimal E305-loop flexibility in human GS, it is perhaps unsurprising that a large post-translational modification applied to a regulatory loop of GSIB could have a dramatic functional impact. The E305-loop thus is likely a hotspot for GS regulation via multiple distinct routes across evolution.

Here, we observe a connection between the filamentation and E305-loop flexibility. There is evidence that GS homologs from bacteria, archaea, and eukaryotes also form a filamentous, head-on-head structure, but their structural and functional consequences are incompletely understood. In E. coli, GS filaments could be formed from supraphysiological levels of select divalent cations that are chelated by adjacent helices across repeating dodecamer units. While high structural resolution was achieved, the catalytic and cellular consequences of filament formation were not explored. Furthermore, E. coli filaments dissolved under turnover conditions leaving the functional relevance unclear. By contrast, in S. cerevisiae, GS filaments were demonstrated to negatively regulate GS activity in the cell and shown to be a crucial adaptation upon starvation conditions (Petrovska et al. 2014). Here, we provide a structural rationale describing how filament formation, stabilized by product, can structurally alter GS E305-loop to tune the substrate-limiting Michaelis constant for ammonia. Importantly, point mutations of the interfacial residues do not show a *K*_M_*,_ammonia_*defect in the presence of glutamine, indicative of the importance of the filament form for product feedback. Thus, the glutamine stabilized filament formation model for human GS represents a new mode of product-based allosteric feedback inhibition for this enzyme class.

Human GS is expressed in many tissues to serve different metabolic needs. Translation and subsequent degradation of GS can take multiple hours to achieve robust expression changes (Arad et al. 1976; Van Nguyen et al. 2016, 2017; Van Nguyen 2021; Zhao et al. 2025) which is potentially insufficient to meet the changing metabolic needs of a given cell. Therefore, regulated enzyme activity through filament formation represents a mode of high temporal control over enzyme activity as it is not reliant on transcription, translation, or degradation of enzymes. Additionally, the Michaelis constant for ammonia report herein (57 μM for decameric GS and 170 μM for filamentous GS) corresponds closely to the concentrations of ammonia measured in cells, which suggests that this modulation of GS activity could have an immediate impact on intracellular turnover rates.

Moreover, our observations support the concept that allostery operates through ensemble redistribution rather than through discrete conformational switches. The filament state of human GS does not simply toggle the E305-loop between rigidly defined “open” and “closed” states, but rather shifts the probability distribution across a spectrum of conformations that collectively impact enzyme activity. This is similar to observations in IMPDH2 and PRPS filaments (Hvorecny et al. 2023; Johnson and Kollman 2020), where assembly modulates the conformational landscape accessible to regulatory elements. Both loop dynamics and filamentation are frequently implicated in tuning activity of enzymes; here a product feedback mechanism links these two concepts together. Collectively, the filament formation of human GS described here is a complex manifestation of ensemble-based allostery to enable timely regulation of enzyme activity.

## Methods

| Key Resources Table |  |  |  |  |
| --- | --- | --- | --- | --- |
| Reagent type (species) or resource | Designation | Source or reference | Identifiers | Additional information |
| gene (*Homo sapiens*) | GLUL | UniProt | P15104 |  |
| strain, strain background (*Escherichia coli*) | BL21(DE3) | New England Bioscience | C2527H | Chemical comp cells |
| cell line (*Homo sapiens*) | HEK293E *GLUL*<sup>-/-</sup> | PMID: **29359789** |  |  |
| cell line<br>(*Homo sapiens*) | HEK293E KO-TetBxb1BFP | This study | Clone 15 | Verified by glutamine auxotrophy that was recovered by DOX induction of wild-type GLUL after integration into landing pad |
| cell line<br>(*Homo sapiens*) | HEK293E GLUL-variants (wild-type, C53A, K52A, R298A, L300A, P242stop) | This study | Clone 15 |  |
| recombinant DNA reagent | pDEST hsGS wild-type and associated variants | This paper |  | His<sub>8</sub>-SUMO tagged GLUL variants for E. coli expression |
| recombinant DNA reagent | Intein-CBD-His<sub>6</sub>-GS | This paper |  | His<sub>8</sub>-SUMO tagged GLUL variants for E. coli expression |
| recombinant DNA reagent | pFGET1 – Ulp1 | Addgene #64697 |  | His<sub>6</sub>-Ulp1 protease expression plasmid |
| recombinant DNA reagent | pBxb1 | PMID: 31612958 |  | Bxb1 recombinase expression plasmid |
| recombinant DNA reagent | attB-GLUL-IRES-mCherry | This study |  | Integration and expression plasmid for GLUL. Wildtype and associated variants. |
| chemical compound, drug | BSG (Bis-sulfosuccinimidyl glutarate) | ThermoFisher | 22323 | Amine-to-amine crosslinker |
| chemical compound, drug | BMOE (bis-maleimidoethane) | ThermoFisher | 21610 | Sulfhydryl-to-sulfhydryl crosslinker |
| software, algorithm | CryoSPARC | PMID: 28165473 |  |  |
| software, algorithm | ChimeraX | PMID: 37774136 |  |  |
| software, algorithm | Phenix | PMID: 31588918 |  |  |
| software,<br>algorithm | ISOLDE | PMID: 29872003 |  |  |
| software,<br>algorithm | Coot | PMID: 31730249 |  |  |
| software,<br>algorithm | GROMACS | PMID: 33032406 |  |  |
| software,<br>algorithm | PLUMED | PMID: 31363226 |  |  |

### Expression and purification of scarless human glutamine synthetase

An expression vector for a 6xHIS-conjugated human glutamine synthetase was ordered from ATUM. The vector was transformed into BL21(DE3) *E. coli* cells and grown overnight at 37 °C in Luria broth with 100 μg ml^−1^ carbenicillin. 5 mL of this starter culture was added to 1 L of the same medium and allowed to grow at 37 °C until an OD600 of 0.6-0.8 was reached. At this point, expression of GS was induced with 0.5 mM IPTG and the temperature was reduced to 18 °C for 16 - 18 hours. The cells were harvested by centrifugation at 4500 rpm for 15 minutes at 4 °C. The cell pellets were transferred to a 50-mL conical tube and stored at −80 °C.

For purification, cells were thawed and resuspended in 60 mM HEPES pH 7.6, 50 mM NaCl, 50 mM MgSO_4_, 0.5 mM TCEP, 10% glycerol followed by lysis by sonication. Lysate was clarified by centrifugation and loaded onto HisTrap FF 5 mL (Cytiva) columns using a peristaltic pump, washed with 60 mM HEPES pH 7.6, 50 mM NaCl, 50 mM MgSO_4_, 0.5 mM TCEP, 10% glycerol, 25 mM imidazole, and eluted with 60 mM HEPES pH 7.6, 50 mM NaCl, 50 mM MgSO_4_, 0.5 mM TCEP, 10% glycerol, 250 mM imidazole. Eluate was concentrated before being aliquoted and flash frozen and stored at −80°C. Thawed aliquots were loaded onto a Superose 6 10/300 column preequilibrated with 60 mM HEPES pH 7.6, 50 mM NaCl, 50 mM MgSO_4_, 0.5 mM TCEP and assayed for activity immediately after size-exclusion chromatography.

### Expression and purification of scarless human glutamine synthetase

Scarless human glutamine synthetase was produced by either of two independent methods: N-terminal His-SUMO tagging with Ulp1 cleavage or C-terminal intein-chitin binding domain-His with DTT induced self cleavage. Both purification methods yielded highly functional GS, however, better yields were obtained via the His-SUMO tagging approach.

Human glutamine synthetase was codon optimized for E. coli (ATUM biosciences) and appended with a C-terminal Mxe GyrA intein domain (NEB IMPACT) followed by a chitin-binding domain (CBD) from *Bacillus circulans* (NEB IMPACT) cloned in-frame with a 3′ His_6_-tag and transformed into BL21(DE3) (NEB). Two liters of Terrific Broth (Fisher) were pre-warmed to 37 °C before being inoculated with 10 ml overnight starter culture. Cultures were grown to 0.6 < OD600 < 1.0 and induced with 0.5 mM isopropyl β-D-1-thiogalactopyranoside (GoldBio) and protein expression was allowed to continue for 3 hours. Cells were pelleted at 3,500 r.c.f. for 20 min at 4 °C, resuspended in base buffer (60 mM HEPES pH 7.6, 50 mM NaCl, 50 mM KCl, 10 mM MgCl_2_, 0.5 mM TCEP) supplemented with 2 mg/mL lysozyme, benzonase, and cOmplete™, EDTA-free Protease Inhibitor Cocktail (Roche) and stored at −80°C until purification.

For the purification of GS-intein-CBD-His, cells were thawed and lysed by sonication. Lysate was clarified by centrifugation and loaded onto HisTrap FF 5 mL (Cytiva) columns using a peristaltic pump, washed with base NiA buffer (60 mM HEPES pH 7.6, 50 mM NaCl, 50 mM KCl, 10 mM MgCl_2_, 25 mM imidazole), and eluted with base NiB buffer (60 mM HEPES pH 7.6, 50 mM NaCl, 50 mM KCl, 10 mM MgCl_2_, 250 mM imidazole). Eluates were spiked with DTT to a final concentration 100 mM and stored overnight at 4°C to allow for intein self-cleavage. Cleaved GS was buffer exchanged into NiA buffer via application to a HiTrap 26/10 Desalting column (Cytiva). His_6_-tagged Mxe GyrA intein-CBD was removed through an ortho NiNTA where sample was applied to a HisTrap High Performance 5 mL column (Cytiva) and the flow through fraction was collected, concentrated in an Amicon 30k MWCO centrifugal filter, spiked with glycerol to a final concentration of 10%, aliquoted, and flash frozen for long term storage at −80°C.

Human glutamine synthetase was codon optimized for E. coli (ATUM biosciences) and subcloned into a pET29a vector bearing an inframe 5′ His_6_-SUMO tag and transformed into BL21(DE3). Two liters of Terrific Broth (Fisher) were pre-warmed to 37 °C before being inoculated with 10 ml overnight starter culture. Cultures were grown to 0.6 < OD600 < 1.0 and induced with 0.5 mM isopropyl β-D-1-thiogalactopyranoside (GoldBio) and protein expression was allowed to continue for 3 hours. Cells were pelleted at 3,500 r.c.f. for 20 min at 4 °C, resuspended in base buffer (60 mM HEPES pH 7.6, 50 mM NaCl, 50 mM KCl, 10 mM MgCl_2_, 0.5 mM TCEP) supplemented with 2 mg/mL lysozyme, benzonase, and cOmplete™, EDTA-free Protease Inhibitor Cocktail (Roche) and stored at −80°C until purification.

For the purification of His_6_-SUMO-GS, cells were thawed and lysed by sonication. Lysate was clarified by centrifugation and loaded onto HisTrap FF 5 mL (Cytiva) columns using an AKTA start, washed with base NiA buffer (60 mM HEPES pH 7.6, 50 mM NaCl, 50 mM KCl, 10 mM MgCl^2^, 25 mM imidazole; 250 mL), and eluted with base NiB buffer (60 mM HEPES pH 7.6, 50 mM NaCl, 50 mM KCl, 10 mM MgCl^2^, 250 mM imidazole; 50 mL).

Eluates were spiked with Ulp1 (see purification below) and dialyzed overnight at 4°C against base buffer. Cleaved GS was separated from His_6_-tagged His-SUMO through an ortho NiNTA where sample was applied to a HisTrap High Performance 5 mL column (Cytiva) and where scarless GS could elute with application of base buffer with 25 mM imidazole. Scarless GS was concentrated in an Amicon 30k MWCO centrifugal filter, spiked with glycerol to a final concentration of 10%, aliquoted, and flash frozen for long term storage at −80°C.

Prior to experimentation, a scarless GS aliquot was thawed, filtered through a 0.22 μm filter, and applied to a Superose 6 increase 10/300 size-exclusion column pre-equilibrated with base buffer. Unless otherwise stated, the decameric peak corresponding to a 16 mL retention volume was collected and stored at 4°C throughout the duration of experimentation. Activity of this peak was unchanged for up to 2 weeks and all biochemical experiments were performed immediately upon purification.

### Purification of Ulp1 Sumo protease

The expression vector encoding His_6_-Ulp1 protease was a gift from the Verba lab at UCSF obtained originally from Addgene 64697. Ulp1 was expressed and purified as previously described (Guerrero et al. 2015) with minor adaptations detailed below. Briefly, 2L of Terrific broth were pre warmed to 37°C before inoculation with 10 mL of a starter culture of BL21(DE3) cells bearing the His_6_-Ulp1 expression vector. Cells were grown to 0.6 < OD_600_ < 0.8 before temperature was dropped to 20°C and expression was induced with 0.5 mM IPTG overnight. Cells were pelleted at 3,500 r.c.f. for 15 min at 4°C and stored in a 50 mL tube at −80°C until preparation.

Cells were resuspended in Lysis buffer (50 mM HEPES pH 8, 350 mM NaCl, 0.5 mM TCEP, 10 mM imidazole, 2 mg/mL lysozyme, benzonase, and cOmplete™ EDTA-free Protease Inhibitor Cocktail (Roche). Lysate was clarified by centrifugation and loaded onto HisTrap FF 5 mL (Cytiva) columns using an AKTA start, washed with 25 mL of Lysis buffer followed by 25 mL of Buffer A (50 mM HEPES pH 8, 350 mM NaCl, 0.5 mM TCEP, 10 mM imidazole) and 25mL of Buffer B (50 mM HEPES pH 8, 350 mM NaCl, 0.5 mM TCEP, 30 mM imidazole). Finally Ulp1 was eluted with 25 mL of Buffer C (50 mM HEPES pH 8, 200 mM NaCl, 0.5 mM TCEP, 300 mM imidazole). Pure fractions were combined and dialyzed against 50 mM HEPES pH 8, 200 mM NaCl, 10 mM EDTA, 0.5 mM TCEP, 20% glycerol). Ulp1 was concentrated to 2 mg/mL and flash frozen in 250 μL aliquots and stored at −80°C.

### Negative stain microscopy

GS was prepared to ∼0.01 - 0.05 mg/mL in base buffer prior to application to glow-discharged carbon-coated grids, stained with 0.7% uranyl formate and imaged on a T12 (FEI) operating at 120 kV. Images were acquired at ×57,000 magnification on a Gatan Rio camera. Two-dimensional classification and averaging was conducted using cisTEM.

### BSG and BMOE crosslinking of GS filament interface

BMOE (bismaleimidoethane; ThermoFisher) and BSG (bis(sulfosuccinimidyl) glutarate-d0; ThermoFisher) powder was thawed to room temperature and reconstituted in DMF to a concentration of 50 mM, aliquoted, flash frozen in liquid nitrogen, and stored at −80°C. Protein samples were diluted to concentrations noted in base buffer (60 mM HEPES pH 7.6, 50 mM NaCl, 50 mM KCl, 10 mM MgCl_2_, 0.1 mM TCEP) and reacted with crosslinker to a final concentration of 0.5 mM for 10 mins at room temperature followed by quench in 5X SDS-PAGE sample buffer (225 mM Tris pH 6.8, 50% glycerol, 0.05 % SDS, 0.2 mg/mL bromophenol blue, 1M DTT) supplemented with 100 mM NH_4_Cl to quench BSG reactions only. Protein concentrations were normalized after quench prior to SDS-PAGE analysis.

### Michaelis-Menten analysis using coupled ATPase assay

ATP-hydrolysis rates were determined using an NADH-coupled assay (pyruvate kinase and lactate dehydrogenase; Sigma) as described previously (Shapiro and Stadtman 1970). Stocks of ATP, NADH, and phosphoenolpyruvate were made in base buffer (60 mM HEPES pH 7.6, 50 mM NaCl, 50 mM KCl, 10 mM MgCl_2_, 0.5 mM TCEP) and the pH was adjusted until it reached 7.5 on ice prior to aliquoting, flash freezing, and storage at −80°C in the dark. NADH was only exposed to light upon thawing and assay set up and no appreciable change in absorbance of control experiments were noted. Briefly, GS was diluted into ATPase mix (final concentrations: 1 mM NADH, 7.5 mM phosphoenolpyruvate, 6 U/mL pyruvate kinase, and 6 U/mL lactate dehydrogenase), applied to a 384-clear bottom plate (Corning) in 10 μL final volume. Steady-state depletion of NADH was assessed by measuring the absorbance at 340 nm in a Spark 10M multimode plate reader (Tecan). Single substrate concentrations were varied under saturating conditions of other substrates as determined by primarily kinetic assessment in at least technical triplicate. Solution pathlength was manually determined per experiment through titration of NADH and used to calculate ATPase rate. Resultant enzyme normalized velocities were plotted against variable substrate concentration and fit using Equation 1:

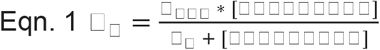

The low concentrations of ammonium required to assess *K*_M,_ *_ammonia_* (< 10 μM) resulted in small changes in total NADH concentration (<1%) relatively quickly and therefore use of full progress curve analysis was pursued as described (Johnson 2019), which captures the rate of substrate depletion or production formation over time as [S] decreases by fitting to the differential equations inherent to the Michaelis-Menten kinetic model. Here, saturating glutamate and ATP concent ations were used in combination with three starting ammonia concentrations (750 μM, 375 μM, and 188 μM) and the NADH-coupled assay components as above in 2X concentration before being mixed with 2X enzyme (0.1 mg/mL final) and the resultant 10 μL reactions (n = 8) were monitored for 5 mins at room temperature in a Spark 10M multimode plate reader (Tecan). Replicates were averaged and the average standard deviation per time point was imported in Kintek Explorer. A standard Michaelis-Menten model was applied to the simulation defined by *k_1_, k_-1_, k_2_, k_-2_,* and *k_3_* as defined in Equation 4.

Scheme 1.

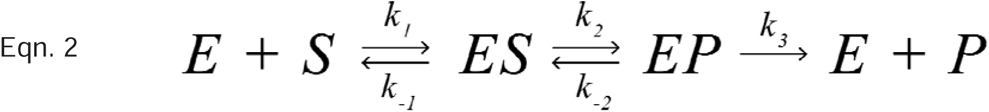

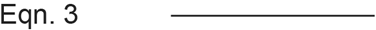

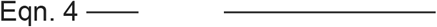

Derived elementary step parameters and their associated uncertainties were used to determine the steady state kinetic parameters with error propagated through each step. The derived *k_cat_* parameters from this procedure were in agreement with those derived under the traditional initial rate procedure.

### Sample preparation for cryoEM analysis

Frozen, mostly pure, GS aliquots were thawed and further purified by size-exclusion chromatography on a superose 6 10/300 column where fractions corresponding to the decamer oligomeric state (∼16 mL) were collected, quantified by A_280_ and tested for activity. Grids (Quantifoil 1.2/1.3 Cu 300 (with or without +2nm C) or Quantifoil 2/2 Au 200) were glow discharged in a Pelco-Easi Glo (15 mA, 10-15s on, 10-15s off) before sample application. Enzyme (0.3-1.3 mg/mL final) in apo (buffer added) or turnover (5 mM ATP, 25 mM ammonium chloride, and 50 mM glutamate; reacted for 53-600s and within steady-state kinetic regime except for the turnover-filament dataset where near full conversion of substrate to product was expected) or ATP bound (5 mM ATP) at room temperature before application to grid held at 4° or 18°C in a Mark IV Vitrobot (ThermoFisher) prior to vitrification. For turnover conditions, all data were collected on Quantifoil 1.2/1.3 Cu 300 +2nm C grids where reactions were initiated at room temperature immediately before being applied to grid in a Vitrobot Mark IV (Thermo) pre-equilibrated to 18°C with 100% humidity and allowed to adsorb to ultrathin carbon layer for 30s before blotting (blot force 3, 11s, Whatman Filter Paper #1) and plunge frozen into liquid ethane. For apo- and ATP-filament datasets, grids were held at 4°C with 100% humidity, blotted 1-3 times with sample, before plunge frozen in liquid ethane. Observation of short filamentous structures was observed with and without depositing 2nm continuous carbon film on grid and preparations

### CryoEM data collection

Wild-type decamer turnover, time-resolved datasets, and R298A decamer turnover GS sample grids were loaded onto a Titan Krios G3i 300 keV microscope (TEM Gamma; SLAC S2C2 Facility, Menlo Park, CA) with a BioQuantum energy filter (20 kV) and Gatan K3 Ceta-D direct detector operating at a nominal magnification of 105,000x corresponding to a physical pixel size of 0.86 Å. A zero-loss peak optimization and gain reference was taken ahead of each data collection. Each dataset was set to a total accumulated dose rate of 40 e-/Å^2^ per micrograph. Each movie was split into 75 frames and a nominal defocus range was set between −0.8 to −2.0 μm. Fringe-free illumination was utilized and image shift was used to collect multiple images within a single hole and beam tilt was adjusted to achieve coma-free alignment during image shift. Automated data acquisition was performed with EPU software.

The low resolution turnover filament dataset was collected at SLAC S2C2 Facility (Menlo Park, CA) on a Talos Arctica G2 200 keV microscope with a Gatan K3 direct detector operating at a nominal magnification of 106,000X corresponding to a physical pixel size of 0.86 Å. A zero-loss peak optimization and gain reference was taken ahead of each data collection. The dataset was set to a total accumulated dose rate of 40 e-/Å^2^ per micrograph. Each movie was split into 40 frames and a nominal defocus range was set between −0.8 to −2.0 μm. Automated data acquisition was performed with EPU software.

The turnover filament dataset was collected at the UCSF CryoEM Core facility where grids were loaded onto a Titan Krios G2 300 keV microscope (Krios 1) with a 20kV energy filter and Gatan K3 direct detector operating at a nominal magnification of 106,000x corresponding to a physical pixel size of 0.82 Å. A zero-loss peak optimization and gain reference was taken by facility management each week. Total accumulated dose rate of 45 e-/Å^2^ per micrograph. Each movie was split into 75 frames and a nominal defocus range was set between −0.8 to −2.0 μm. Fringe-free illumination was utilized and image shift was used to collect multiple images within a single hole and beam tilt was adjusted to achieve coma-free alignment during image shift. Automated data acquisition was performed with SerialEM software.

Apo- and ATP-Filament datasets were collected at the Arnold & Mabel Beckman center for cryo-electron microscopy at the University of Washington where grids were loaded onto a Titan Krios G3 300 keV microscope (Krios 1) with a 20kV Gatan BioQuantum energy filter and Gatan K3 direct detector operating at a nominal magnification of 105,000x corresponding to a physical pixel size of 0.843 Å. Total accumulated dose rate between 60 and 64.2 e-/Å^2^ per micrograph.

### CryoEM image processing and model building

Movies were aligned, motion-corrected, and dose-weighted using Patch Motion Correction in cryoSPARC (v3.1 and above) or cryoSPARC Live, except for the turnover filament dataset which was motion-corrected and dose-weighted on-the-fly with MotionCor2 (v1.3.0). Contrast transfer function (CTF) estimation was performed in cryoSPARC using the Patch CTF except for the turnover filament dataset which was estimated using CTFFIND4 ‘on-the-fly’ as part of the Scipion (v2.0) software suite. Micrographs with poor CTF fit (generally >5 Å and/or containing crystalline ice) were discarded. All further processing steps were performed in cryoSPARC (v3.1 or higher) where blob picked particles were classified in two dimensions and non-background particles were subject to either further rounds of template picking and 2D classification or used for 3D homogeneous refinement using either *ab initio* prepared input maps or using maps from other datasets as 3D refinement input. Where applicable, particles were further selected with either or a combination of heterogeneous refinement and/or 3D classification as shown in the supplemental figures. Homogeneous or non-uniform refinement was utilized for final refinement and reconstructions and typically included a symmetry operator (D5 for filament reconstructions, C5 for R298A decamer turnover reconstruction and C1 for wild-type decamer turnover reconstruction). Local map resolution was estimated in cryoSPARC for the final maps. Focused masks were generated in ChimeraX (v.1.7 and above). Focused refinement and 3D classification (3 Å filter resolution, PCA initialization) were performed in cryoSPARC. Strategy of class picking and refinement are noted in**Supplementary Figure 19** and **Supplementary Figure 18**

An initial model, PDB 2OJW, was used to build the apo-filament model and the apo-filament model was used as the initial model for all subsequent model building. Ligands were placed with ISOLDE (v1.7) and those with >0.5 Qscore and supporting biochemical and/or literature precedent were built. Models were built iteratively using the ISOLDE (v1.7) plugin for ChimeraX (v.1.7 and above), Coot (v0.9.8.7), and Phenix (v1.20.1; Real Space Refinement and CryoEM Comprehensive Validation). Models were additionally validated with Q-score as a plugin function for ChimeraX (**Supplementary Table 1**). Final cryoEM maps were sharpened in Phenix using the Autosharpen feature by half-maps.

Decamer:Decamer rotation across the filament interface was determined through inspection of the apo-filament cryoEM map in ChimeraX where an outline of a pentamer from one decameric unit was rotated with respect to the outline of a pentamer across the filament interface and the rotation was depicted in Figure 1D.

### MD-based cryoEM ensemble refinement

Initial models of glutamine synthetase under turnover conditions in the filament (complexed with ATP and MG) and decamer form (complexed with ADP, ATP and MG), as well as a model of glutamine synthetase in the apo-filament form generated from cryoEM data were processed using the CHARMM-GUI server to initialize molecular dynamics simulations (Jo et al. 2008). Each system was solvated in a triclinic box such that the edge of the box was 1.0 nm away from the closest atom of each glutamine synthetase model. K^+^ and Cl^−^ salts were added to the system at a concentration of 0.15 M, along with additional ions to neutralize the charge of each system. The CHARMM36m forcefield was used for the proteins and the CHARMM General Forcefield (CGenFF) (Vanommeslaeghe et al. 2010) was used for the ATP and ADP molecules present in the turnover condition systems. Water molecules were modeled using the mTIP3P model (MacKerell et al. 1998). All simulations were carried out using a leap-frog algorithm with a 2 fs timestep. The smooth particle mesh Ewald method (Essmann et al. 1995) was used for calculating electrostatic interactions within a cutoff distance of 1.2 nm. Van der Waals interactions were gently decreased at 1.0 nm and cutoff at 1.2 nm. All simulations were performed using GROMACS v 2021.5 (Abraham et al. 2015) paired PLUMED v 2.9 (Tribello et al. 2014).

### Details of EMMIVox Simulations

Refined single-structure models and structural ensembles of the GS in the apo-filament, turnover filament, and turnover decamer forms were generated using EMMIVox (Hoff et al. 2024). In brief, EMMIVox is a Bayesian inference approach to determine single-structure models and structural ensembles from 3D cryoEM maps. The direct utilization of experimental cryoEM density and 3D half-maps in the Bayesian framework of EMMIVox automatically balances experimental information with accurate physico-chemical models. Both single structure refinement and ensemble modelling followed the standard EMMIVox protocol (Hoff et al. 2024). CryoEM maps for each GS system and their associated half-maps were pre-processed prior to EMMIVox simulations. Correlated voxels within the experimental 3D maps were removed. This was achieved by first selecting all voxels within 3.5 Å of the model and removing correlated data points with a correlation greater than a pearson coefficient threshold of 0.8. Next, the difference between voxel density for the two half-maps of each system were used to generate a lower bound for the error parameter associated with each density voxel in the EMMIVox framework. Equilibration and production simulations were performed as follows. First, each system was energy minimized using the steepest gradient descent approach. Next, a 10 ns long NPT equilibration was performed using the Bussi-Donadio-Parrinello thermostat (Bussi et al. 2007) and Berendsen barostat (Berendsen et al. 1984) set at 300K and 1 atm respectively. Next, 2 ns long NVT simulations were formed using the same thermostat at 300K. Positional restraints were applied to the NPT and NVT equilibration simulations. Next, an optimal scaling factor between the predicted and observed cryoEM maps was determined using the NVT equilibration simulation trajectory and the PLUMED *driver* tool. Scaling factor values between 0.5 and 1.8 were scanned at 0.05 intervals, with the scaling factor resulting in the lowest EMMIVox hybrid energy score selected for use in the production simulations. B-factors for each chain in the GS were optimized independently without the use of symmetry constraints.

Single-structure refinement production simulations were performed in the NVT ensemble using the thermostat conditions of the NVT equilibration run. Each GS system studied was simulated for 10 ns, with B-factor values sampled every 500 MD steps and a maximum MC move of 0.05 nm^2^. To optimize performance, the EMMIVox hybrid energy was calculated every 4 MD timesteps. After single-structure refinement, the conformation with the lowest EMMIVox hybrid energy and the associated B-factor values were extracted. This structure was then energy minimized while the B-factor values were simultaneously sampled using a modified MC sampler allowing only downhill moves. Any discontinuities in the final structure obtained from the energy minimization due to the PBC were then fixed using the PLUMED *driver* tool. We then generated a PDB of the final refined structure and optimized B-factor values. These models represent the final refined single-structure models of each GS system.

EMMIVox ensemble simulations were performed using the same settings as in the single-structure refinement, with the exception of the B-factors, which were fixed, for all residues, at the lowest value obtained from single-structure refinement as a measure of background noise. 20 metainference replicas were used for each system, with each replica run for 20 ns, resulting in 400 ns of total simulation time per system.

### Pairwise residue distance distributions

The probability distribution of residue-residue distances for selected residue pairs were calculated by taking the minimum distance between any two atoms within the selected residue pair for each conformation present in the structural ensemble extracted from EMMIVox. The calculation was performed for each chain within GS, resulting in 10 residue-residue distances extracted for each conformation in the structural ensemble. Kernel density estimation plots were done using a bandwidth adjustment value equal to 1. Histograms plots were calculated using 150 bins.

### Glutamine synthetase knockout HEK293e cell culture and landing pad clone generation

HEK293E glutamine synthetase knockout cells (Yu et al. 2018) and any derivatives were cultured in Dulbecco’s modified Eagle’s medium (DMEM; high glucose, pyruvate, with glutamine; Gibco) supplemented with 10% dialyzed fetal bovine serum (Gibco), 1x GSEM supplement (Sigma-Aldrich), and Penicillin-Streptomycin (10U Penicillin/10 μg ml^−1^; Gibco). Cells were grown on plates at 37 °C with 5% CO_2_. For routine passaging, cells were detached using trypsin–EDTA 0.25% (Gibco), resuspended in medium, centrifuged for 5 minutes at 200 x g, resuspended and plated.

Landing pad clones were generated as described previously using dAAVS1-TetBxb1BFP (Matreyek et al. 2017, 2020). Successful clones were verified by genomic PCR and co-transfection of attB-EGFP and attB-mCherry. Clone 15 passed all validation tests and was used for all following experiments (referred to in Methods as GS-KO-TetBxb1BFP and *HEK293E GLUL^−/−^* in main text). Long-term passaging of GS-KO-TetBxb1BFP was done with the addition of 2 μg ml^−1^ doxycycline (Sigma-Aldrich).

### Recombination of single-variant clones or libraries into the GS-knockout-landing pad cell line

Single-variant clones were recombined into the Tet-on landing pad in GS-KO-TetBxb1BFP cell. Transfection of attB-GS variant 0.5 × 10^6^ cells per well transfected with 1.5 μg attB-plasmid DNA and 1.5 μg pCAG-HA-NLS-coBxb1 using Lipofectamine 3000 (Thermo) per manufacturer recommendations. Two days post-transfection, cell media was refreshed with the addition of 2 μg ml^−1^ doxycycline. Four days post-transfection, cells were selected with media containing 2 μg ml^−1^ doxycycline (Sigma-Aldrich) and 0.25 μg ml^−1^ puromycin (Gibco). Cells were selected for one week under puromycin and the resultant stable cell lines were frozen in 10% DMSO (Sigma) in FBS (Gibco) for future experiments.

### Growth test of wild type GS and disease mutants in GS-KO-TetBxb1BFP

To determine the growth rate of different glutamine synthetase variants in the presence and absence of glutamine, stable *HEK293E GLUL_variant_* cell lines (wild-type, C53A, K52A, R298A, L300A, or P242stop) were plated at low confluence (10,000 cells per well) in a series of identical white, tissue culture treated, 96-well plates (GreinerBio) in DMEM (+glutamine and 10% dialyzed FBS, GSEM, 10U Penicillin, 10 μg ml^−1^ Streptomycin, 2 μg ml^−1^ doxycycline, and 0.25 μg ml^−1^ puromycin) and allowed to adhere to the plate for 4 hours. Subsequently, media was swapped (DMEM with 10% dialyzed FBS, GSEM, 10U Penicillin, 10 μg ml^−1^ Streptomycin, 2 μg ml^−1^ doxycycline, and 0.25 μg ml^−1^ puromycin and with or without glutamine) and a Day ‘0’ cell count time point was measured using the CellTiter-Glo kit (Promega). Following the protocol from the vendor, CellTiter-Glo reagent was mixed 1:1 ratio with cells and incubated for 10 mins at room temperature to lyse and provide a luminescence readout. Luminescence was measured on a Veritas luminometer and converted to cell counts from a standard curve of *HEK293E GLUL_wild-type_*.

## Data availability

Cryo-EM structures and atomic models have been deposited in the Electron Microscopy Data Bank (EMDB) and Protein Data Bank (PDB), respectively, with the following accession codes: EMD-70845, PDB: 9OTQ (Apo-Filament); EMD-70841, PDB: 9OTM (Turnover Filament); EMD-77146 (Focused Refinement Turnover Filament); EMD-70843, PDB: 9OTO (Turnover Decamer); EMD-70842, PDB: 9OTN (ATP bound Filament); EMD-70844, PDB: 9OTP (R298A turnover). Aligned and dose-weighted micrographs used to develop the turnover-filament structure (EMD-70841) are publicly available on EMPIAR (EMPIAR-13621). Initial extracted, template-picked particles from the time-resolved cryo-EM datasets are publically available on EMPIAR (EMPIAR-13622). EMMIVox ensemble refinement and trajectory files are available on Zenodo (10.5281/zenodo.20298855). Maps emerging from focused classification or focused refinement are also available on Zenodo (10.5281/zenodo.20298855). All the GROMACS topologies and PLUMED input files needed to reproduce our EMMIVox calculations as well as the final refined single-structure models are available in PLUMED-NEST (www.plumed-nest.org), the public repository of the PLUMED consortium (PLUMED consortium 2019), as plumID:25.017.

## Materials availability

Plasmids generated in this study are available upon request from the corresponding author (JSF). Cell lines generated in this study (HEK293E KO-TetBxb1BFP and HEK293E GLUL variants) are available upon request and have been cryopreserved for long-term storage.

## Acknowledgements

We acknowledge funding for ERG (F32GM144982),JSF (R35 GM145238), SEH (Pasteur-Roux-Cantarini postdoctoral fellowship), and JMK (R35 GM149542). We acknowledge Douglas Fowler and Kenny Matreyek for providing constructs and advice for landing pad integration. The time-resolved cryoEM datasets that led to the reconstruction of the turnover-decamer and another dataset leading to the R298A turnover decamer reconstruction were collected at the S2C2 Stanford-SLAC CryoEM Center which is supported by the National Institute of General Medical Sciences (1R24GM154186). Cryo-EM equipment at UCSF is partially supported by NIH grants S10OD020054, S10OD021741 and S10OD026881 and Howard Hughes Medical Institute.

---

**Figure 1-figure supplement 1.**
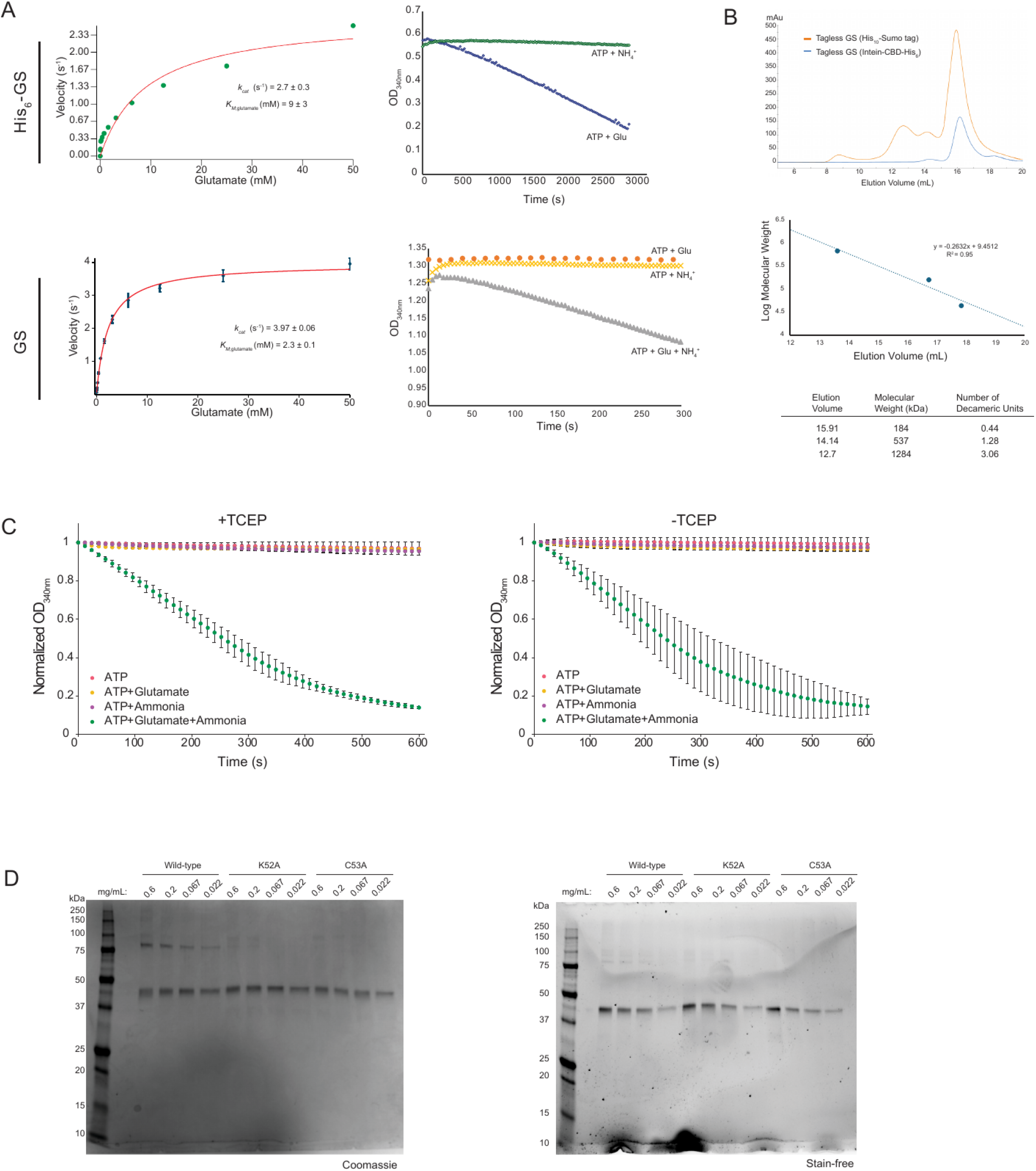
Biochemical characterization of recombinant human GS. A) Steady-state kinetic analysis of N-terminally His_6_-tagged GS (top) and tagless GS derived from the GS-Intein-CBD-His_6_ tagged construct where the *K_M_*_glutamate_ is noted to be 3-fold higher for His_6_-tagged GS and His_6_-tagged GS is noted to display non-specific ATP hydrolysis under ATP + Glutamate conditions suggesting potential loss of γ-glutaryl phosphate intermediate. Kinetic constants were determined from n=3 technical replicates. B) Size-exclusion chromatography traces of tagless GS constructs derived either from the His_10_-Sumo tagged GS or Intein-CBD-His_6_ tagged GS where in both cases the predominant molecular weight species is found at ∼16 mL elution volume on a Superose 6 increase (Cytiva) column which corresponds to the decameric species. Molecular weight calibration curve and predicted molecular weights are included below. C) Steady-state NADH coupled-enzyme assays were performed to validate enzymatic coupling in the presence and absence of TCEP. In all cases, robust ADP production for GS was only observed when all three substrates were present. Kinetic traces were determined from n=3 technical replicates. D) Full SDS-PAGE gel images for bifunctional crosslinking experiments presented in Figure 1E using Bis-sulfosuccinimidyl glutarate (BSG; left) or bis-maleimoethane (BMOE; right).

**Figure 1-figure supplement 2.**
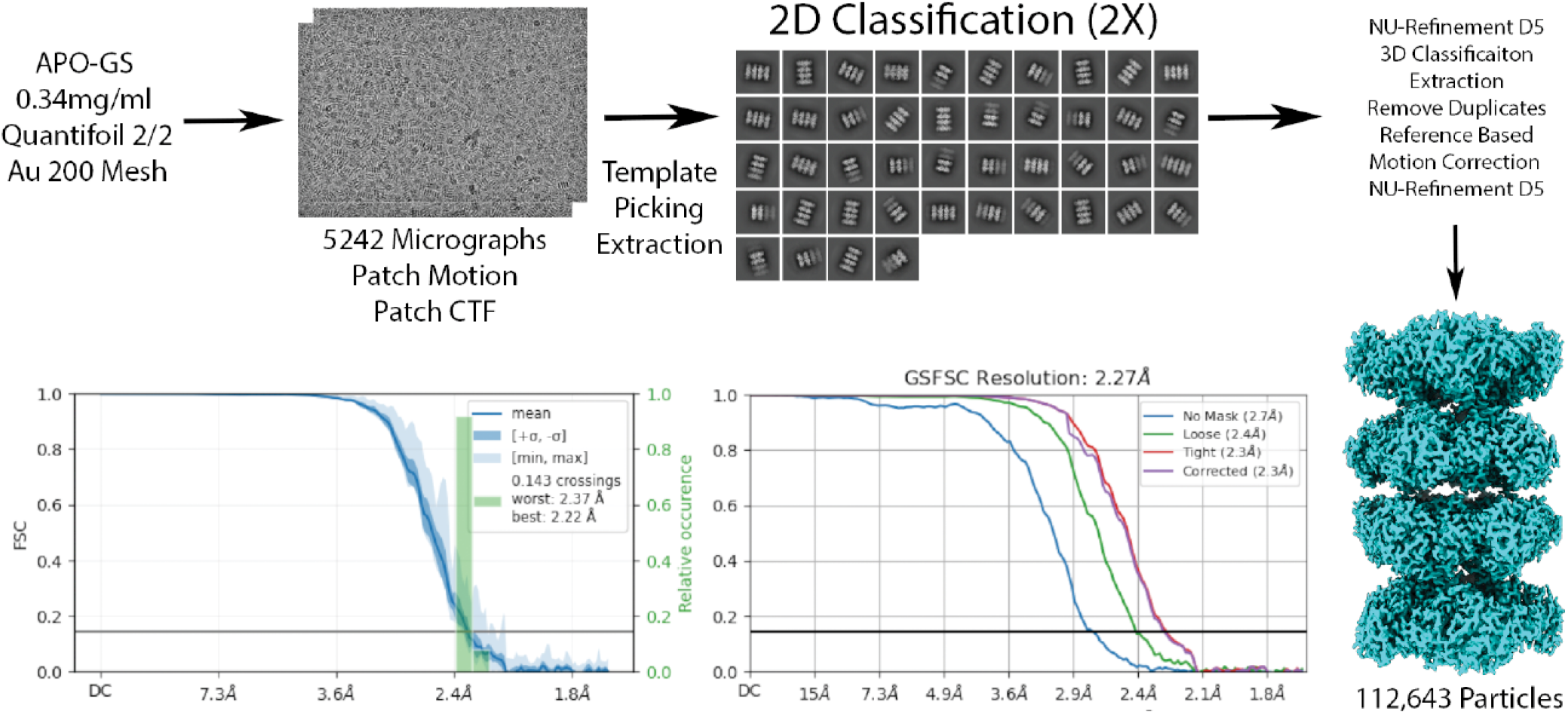
cryoEM data processing pipeline for apo-filament map. Depicted in the pipeline includes grid type, protein concentration, representative micrographs, utilized 2D classes, 3D reconstruction details, and final maps with associated particle counts and FSC curves.

**Figure 1-figure supplement 3.**
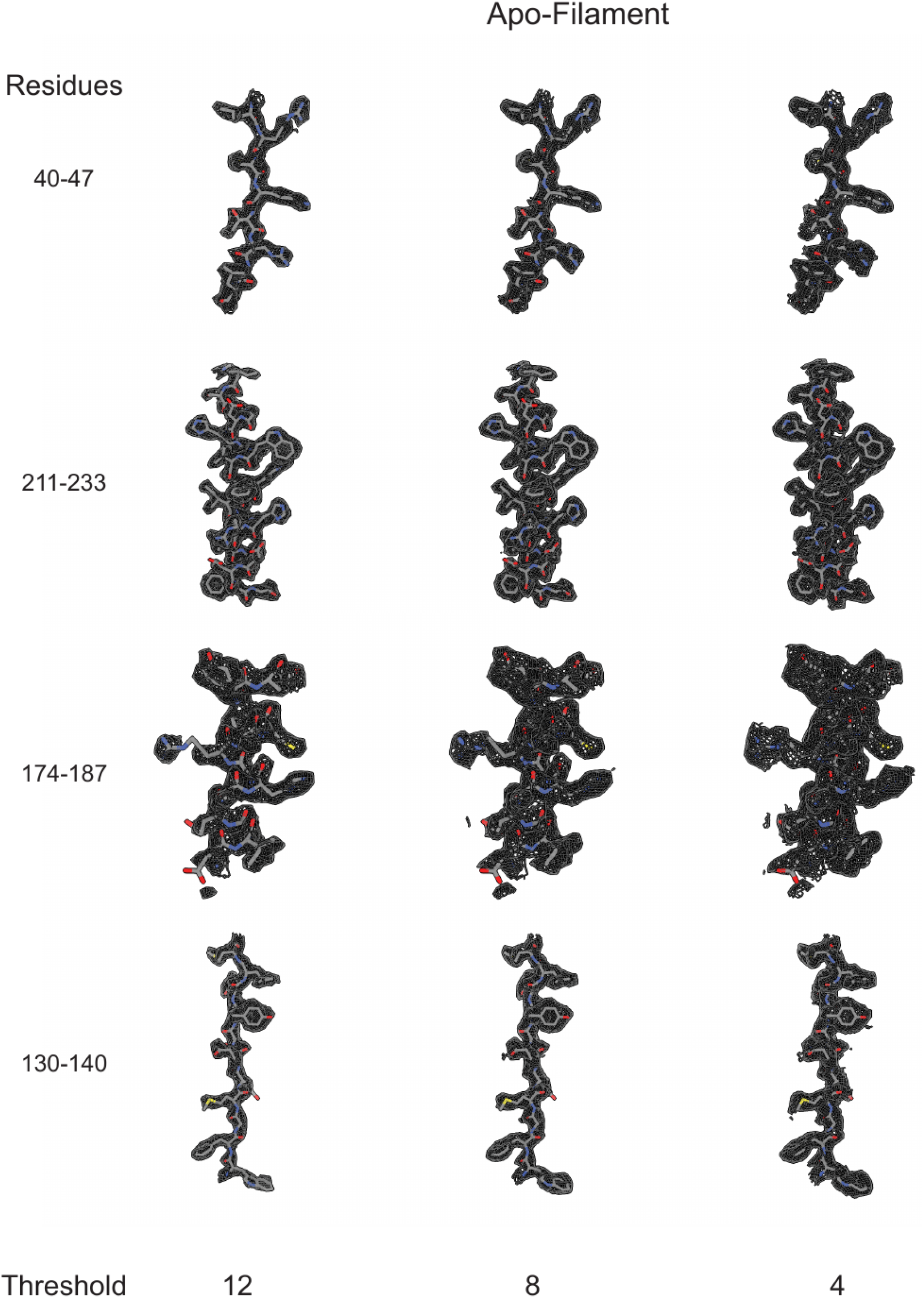
Sharpened cryo-EM density (see Methods) for the apo-filament is presented for four regions of the map containing either β-sheet (residues 40-47 and 130-140) or α-helix (residues 174-187 and 211-233) to demonstrate side chain density and fit.

**Figure 1-figure supplement 4.**
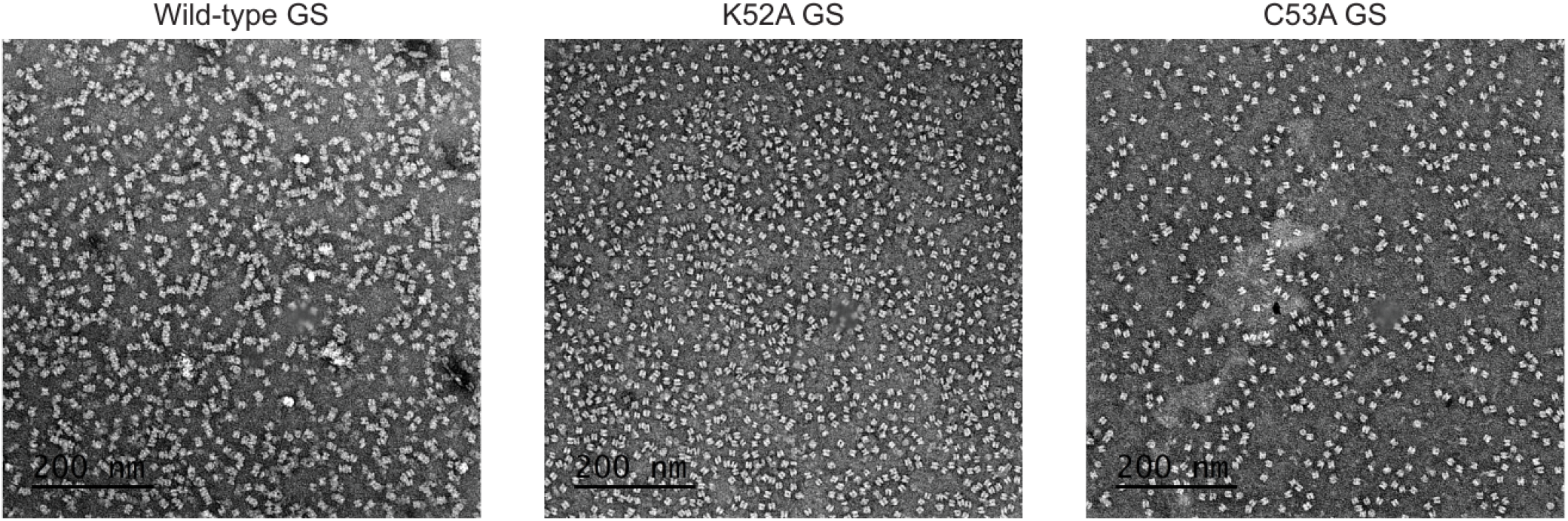
Negative stain EM of human glutamine synthetase variants showing attenuation of filament formation for the point mutants K52A (middle) and C53A (right) compared to wild-type (left).

**Figure 2-figure supplement 1.**
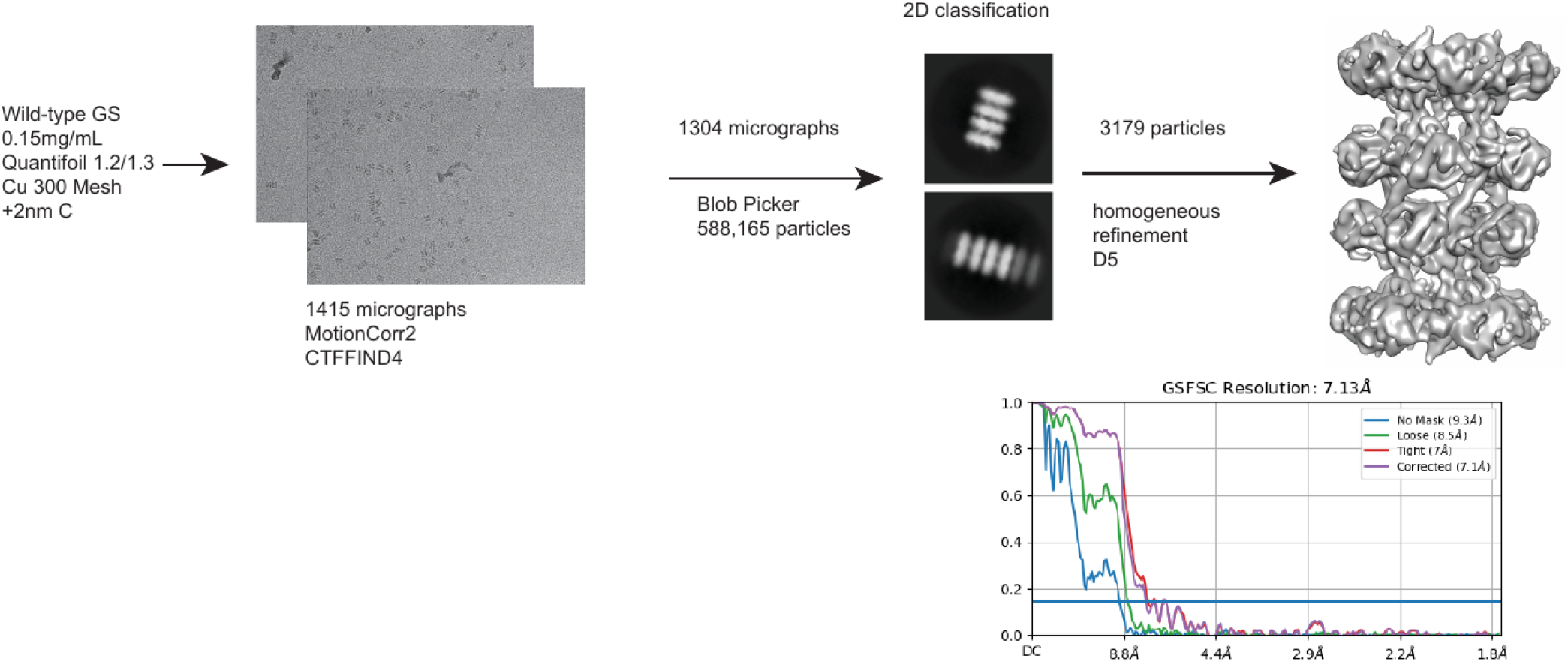
cryoEM data processing pipeline for initial low resolution turnover-filament. cryoEM data processing pipeline for low resolution turnover filament map. Depicted in the pipeline includes grid type, protein concentration, representative micrographs, utilized 2D classes, 3D reconstruction details, and final map with FSC curve.

**Figure 2-figure supplement 2.**
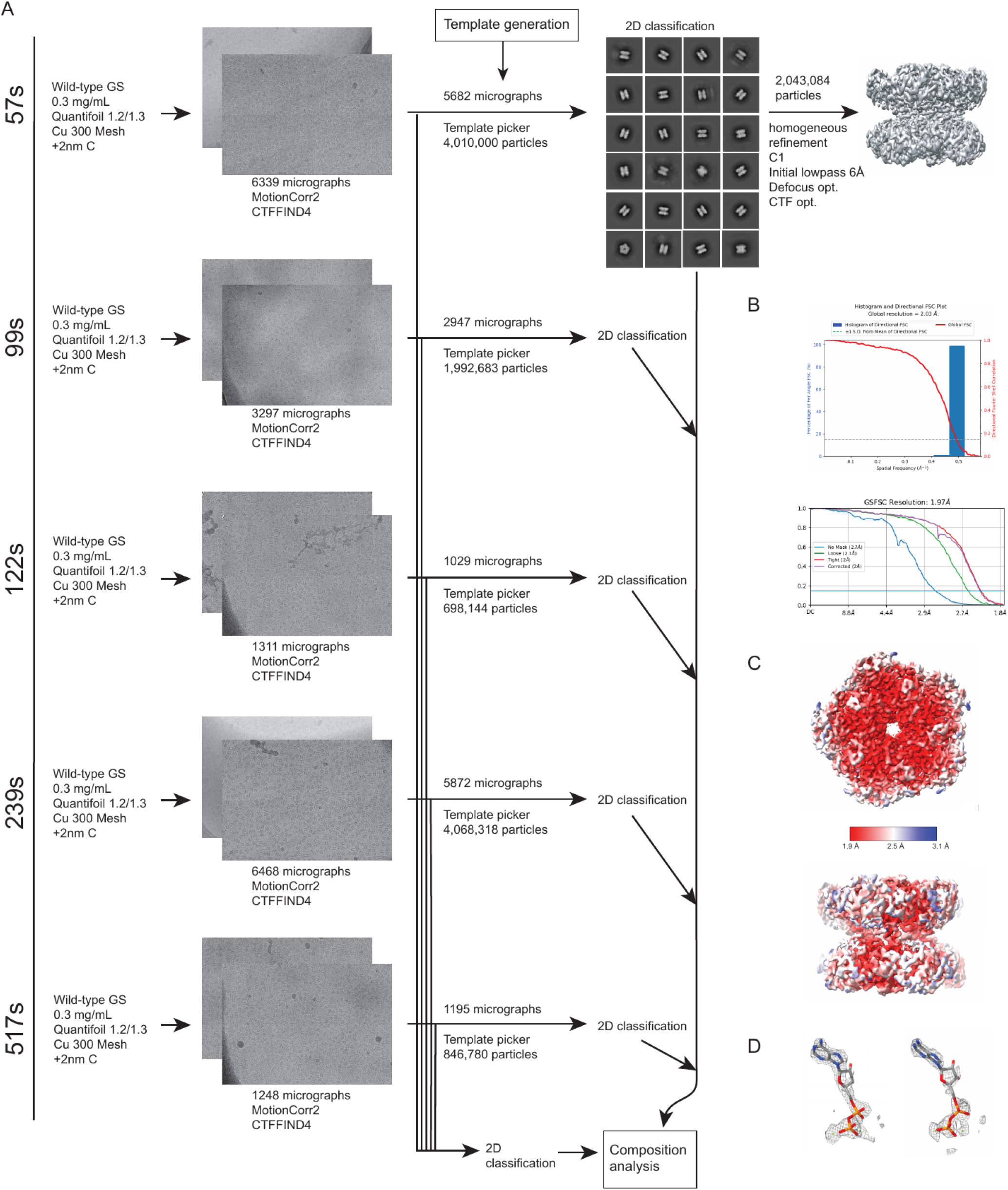
cryoEM data processing pipeline for time-resolved datasets and turnover-decamer map. A) Depicted in the pipeline includes five total datasets where in each dataset, the grid type, protein concentration, representative micrographs, and 2D processing workflow. In addition, the 57s turnover dataset was for 3D reconstruction of the decamer turnover map and displays the utilized 2D classes, 3D reconstruction details for the final map (unsharpened) with associated particle counts. B) FSC curves for the final turnover decamer map. C) Local resolution estimation turnover decamer map. D) Active site ADP and Mg(II) density (sharpened) depiction at threshold 12 for chain A (left) and chain G (right).

**Figure 2-figure supplement 3.**
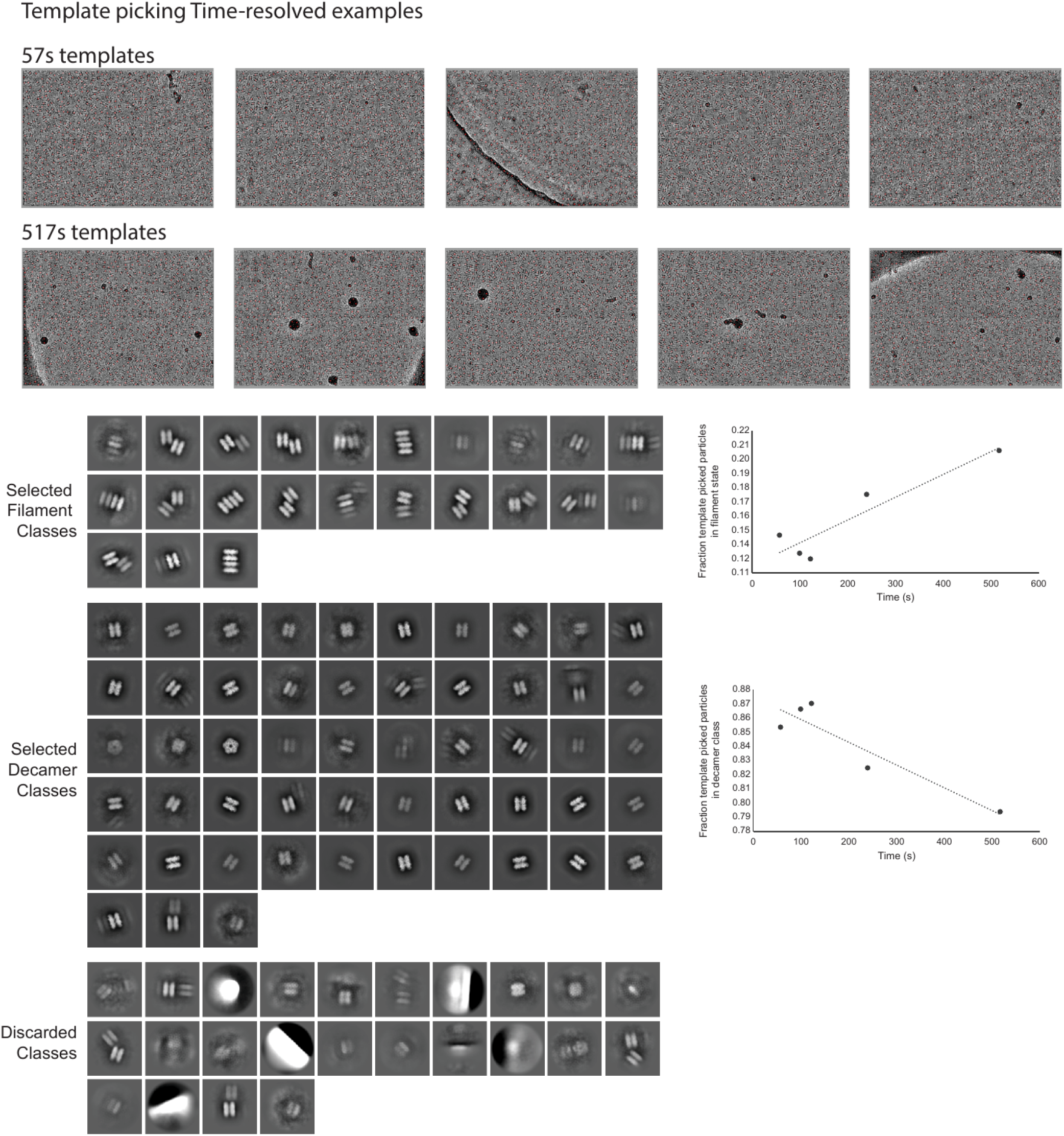
Global 2D classification of time-resolved datasets. Representative template picked particle images for the 57s and 517s datasets are depicted above. All five tr-cryoEM datasets were combined and classified in two dimensions and separated into filament, decamer, and discarded classes with 2D class images depicted at bottom left. 2D classes where partial head-on-head contacts could be discerned were included in the filament classification. 2D classes where multiple decamers were present, but not in a head-on-head arrangement were discarded. The filament and decamer particles classes were then referenced to the original dataset using cryoSPARC tools and the fraction of each species was quantified and linearly fit (bottom right).

**Figure 2-figure supplement 4.**
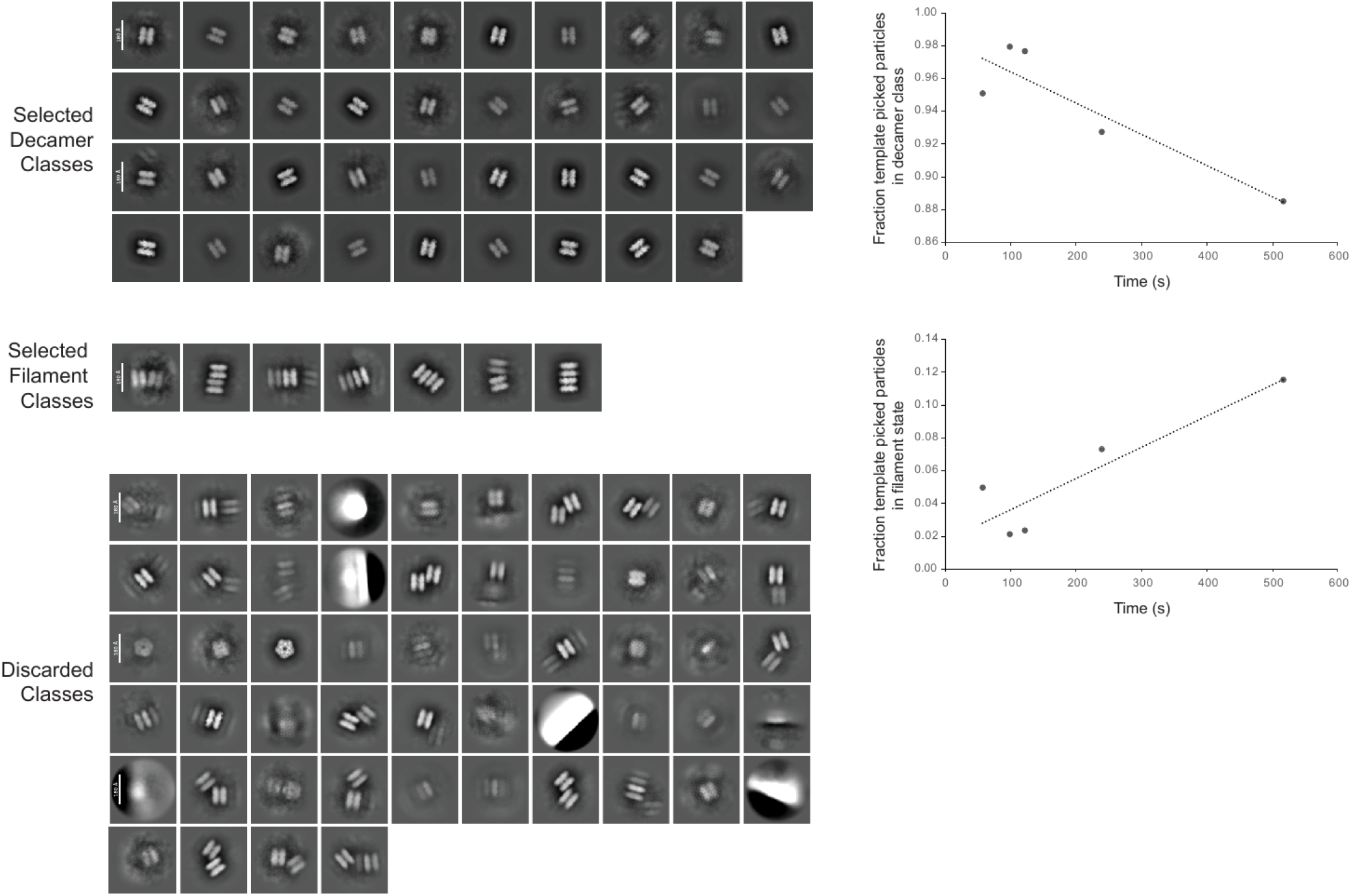
Global 2D classification of time-resolved datasets with more stringent particle picks for quantification. All five tr-cryoEM datasets were combined and classified in two dimensions and separated into filament, decamer, and discarded classes with 2D class images depicted at bottom left. 2D classes where partial head-on-head contacts could be discerned were included in the filament classification. 2D classes where multiple decamers were present, but not in a head-on-head arrangement were discarded. 2D classes representing the ‘top’ view (showing all five pentamer edges) were not quantified as these could not be confidently related to either a filament or decamer class. The filament and decamer particles classes were then referenced to the original dataset using cryoSPARC tools and the fraction of each species was quantified and linearly fit (right).

**Figure 2-figure supplement 5.**
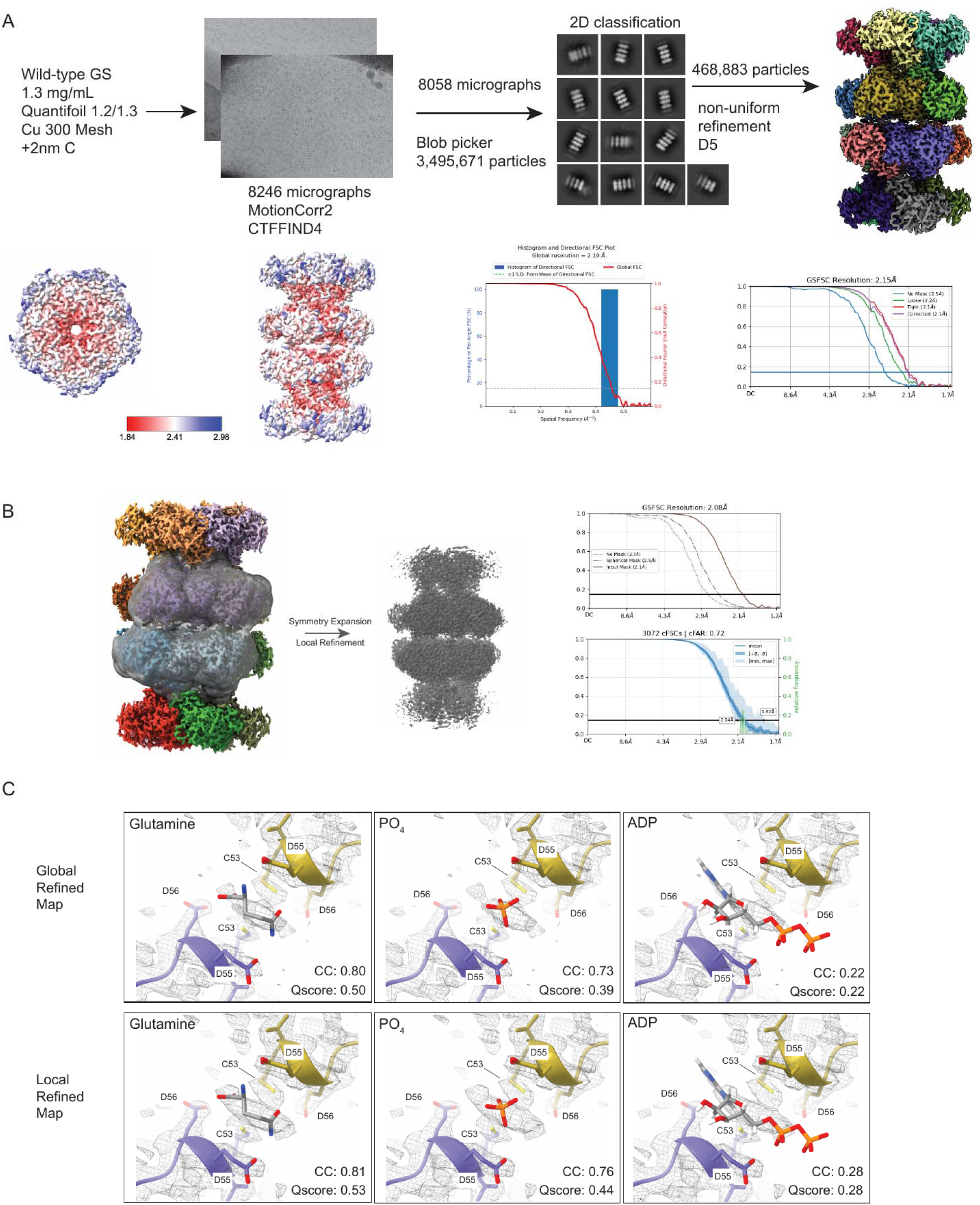
A) Cryo-EM data processing pipeline for high resolution turnover-filament. Depicted in the pipeline includes grid type, protein concentration, representative micrographs, utilized 2D classes, 3D reconstruction details, and final map (sharpened) with associated particle counts. Below are local resolution estimation map (unsharpened) and FSC curves for the final map. B) Local refinement of the filament interface was conducted by symmetry expanding the particles used in the global refinement by their D5 symmetry operator, constructing a mask centered on one glutamine bound filament interface and including portions of four neighboring subunits before local refinement was conducting without symmetry. C) GS products (glutamine, PO_4_, and ADP) individually fit into the filament interface density of the global refined map and for the locally refined map. Each map was sharpened per procedure in Methods and correlation values (CC) and Qscore values are included for each fit.

**Figure 2-figure supplement 6.**
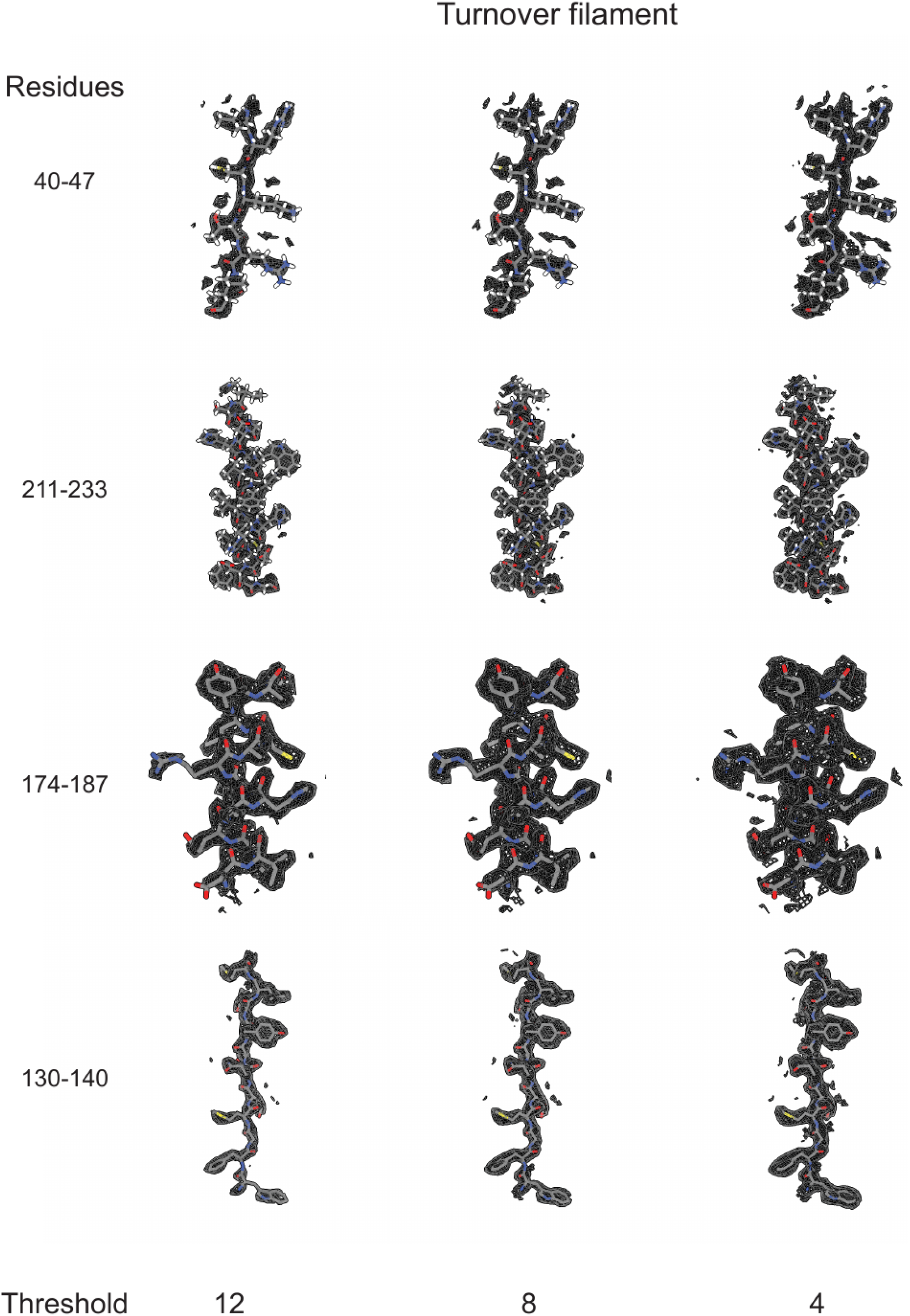
Sharpened cryo-EM density (see Methods) for the turnover filament dataset is presented for four regions of the map containing either β-sheet (residues 40-47 and 130-140) or α-helix (residues 174-187 and 211-233) to demonstrate side chain density and fit.

**Figure 2-figure supplement 7.**
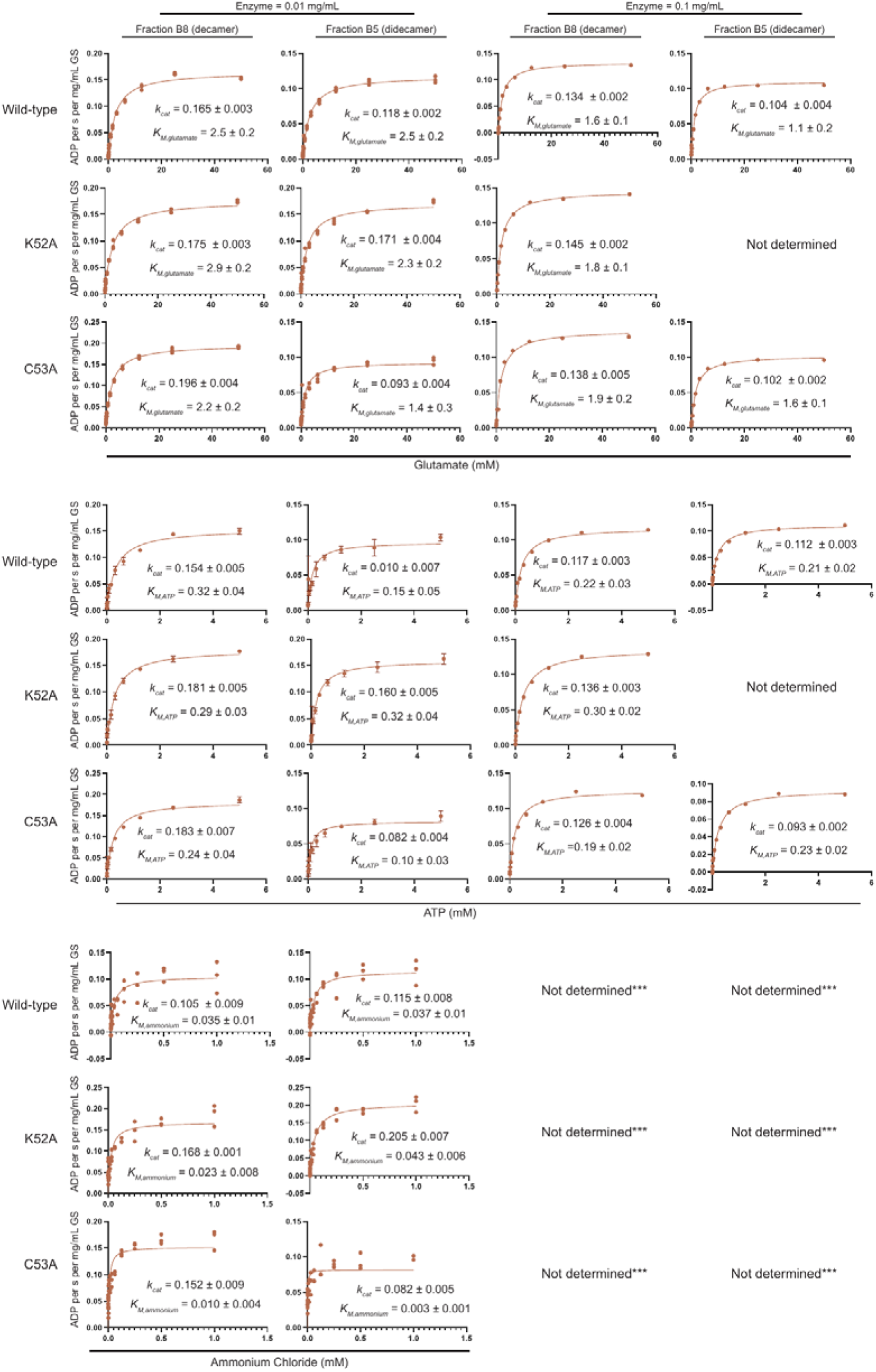
Steady-state kinetic screening with Michaelis-Menten fitting for wild-type, K52A, and C53A GS variants. Initial velocities from the coupled-ATPase assay measurements are presented with fits to a basic Michaelis-Menten model included. Error was estimated from goodness of fit as S.E.M. Enzyme was assayed at 0.01 mg/mL (left) or 0.1 mg/mL (right) and sample from either the B8 (decamer) or B5 (2-decamer) fraction from size-exclusion chromatography from a Sup6 Increase column. The K52A and C53A variants displayed nearly identical Michaelis-Menten parameters as wild-type under both conditions. K52A B5 fraction was not of high enough concentration to assay at 0.1 mg/mL and thus not determined. *** indicates that the rate of conversion at 0.1 mg/mL enzyme was too fast to capture by initial velocity analysis. Kinetic constants were determined from n=1 for 0.1 mg/mL enzyme concentration due to enzyme yield limitations and n=3 technical replicates for 0.01 mg/mL enzyme concentration and were treated as diagnostic regarding oligomeric state influence on kinetic constant profiles.

**Figure 2-figure supplement 8.**
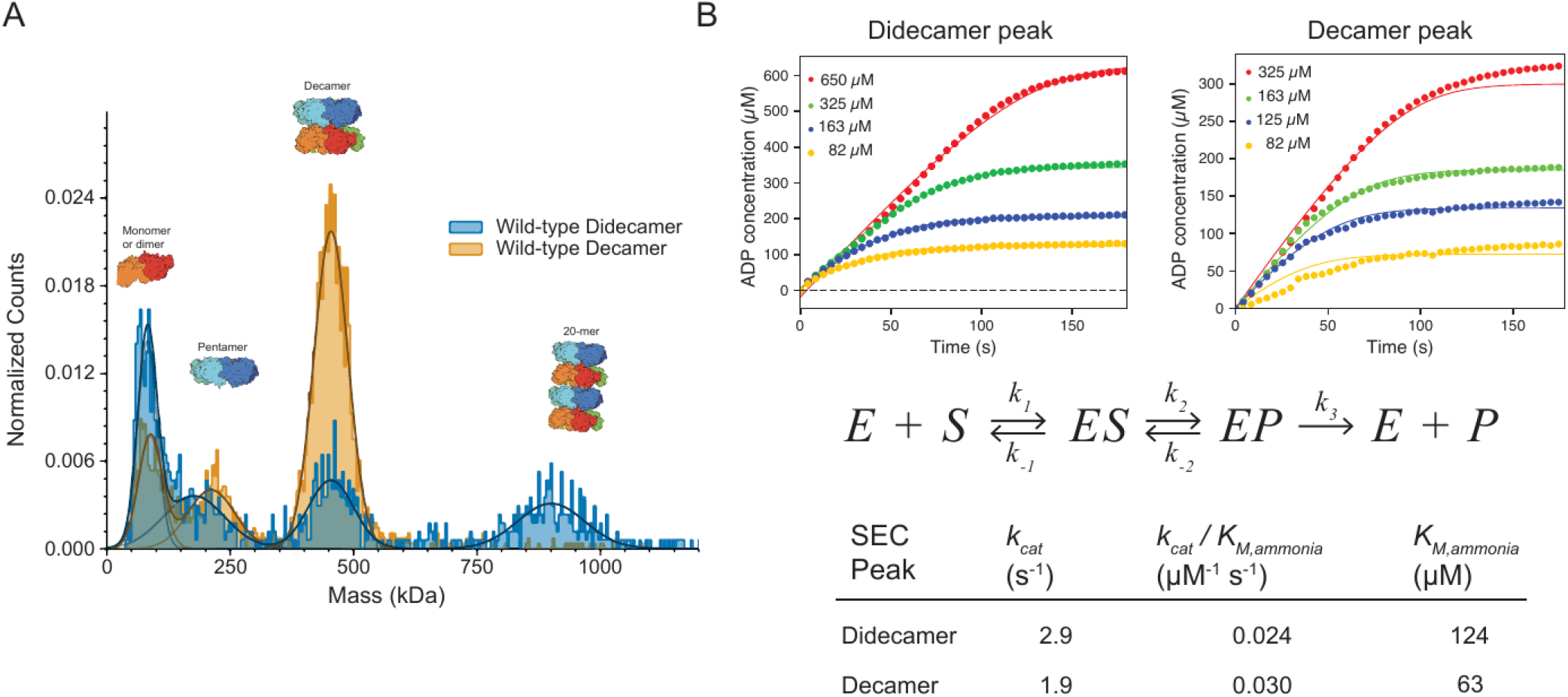
Initial assessment of filament steady-state kinetics. A) mass photometry assessment of oligomeric state for wild-type GS at 0.01 mg/mL demonstrating presence of 2-decameric enzyme in the 2-decameric SEC peak but not in the decameric SEC peak fraction. Additionally observed is the multiple oligomeric states GS could be found in solution at lower concentration. B) Steady-state product formation data for wild-type GS 2-decameric or decameric SEC peaks performed at 0.1 mg/mL GS with varied ammonia concentrations. Top, ADP product concentration (n=1 per concentration) per time at multiple ammonia concentrations with saturating glutamate and ATP globally fit to a simple Michaelis-Menten Model shown in middle using Kintek Explorer. Fit parameters displayed at bottom demonstrate a 2-fold change in *K_M,ammonia_*. An approximate sigma value of 30 μM was used for fitting each dataset.

**Figure 2-figure supplement 9.**
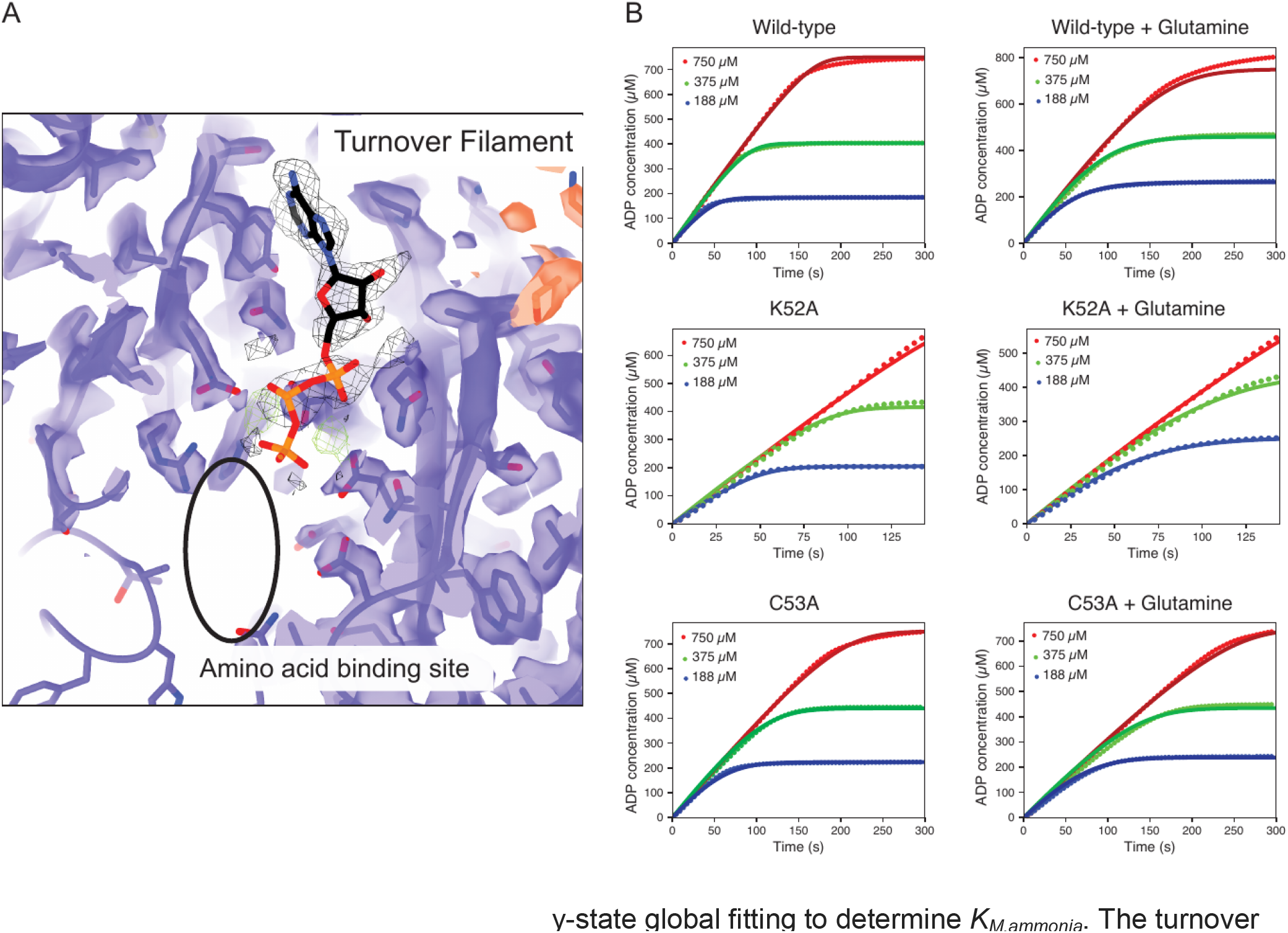
Glutamine-stabilized filament steady-state global fitting to determine *K_M,ammonia_*. The turnover filament map is presented in purple (chain A), black mesh (ligand of chain A), and orange (chain E). Active site residues are shown from the consensus EMMIVox model. Absence of density corresponding to the amino acid binding pocket is shown (black circle) demonstrating that despite high glutamine levels in this dataset, no glutamine is bound to the active site. Right, steady-state data is presented as an average of eight independent progress curve datasets with GS at 0.1 mg/mL, glutamate at 50 mM starting concentration, ATP at 5 mM starting concentration, and ammonium chloride at 0.75, 0.375, and 0.188 mM as indicated. Data were globally fit to a simple Michael-Menten model (bottom left) using Kintek Explorer. Fits are included in plotted data. An average sigma value of 50 μM was used for fitting each dataset. Raw data are included in Figure 2 Source Data 2.

**Figure 3-figure supplement 1.**
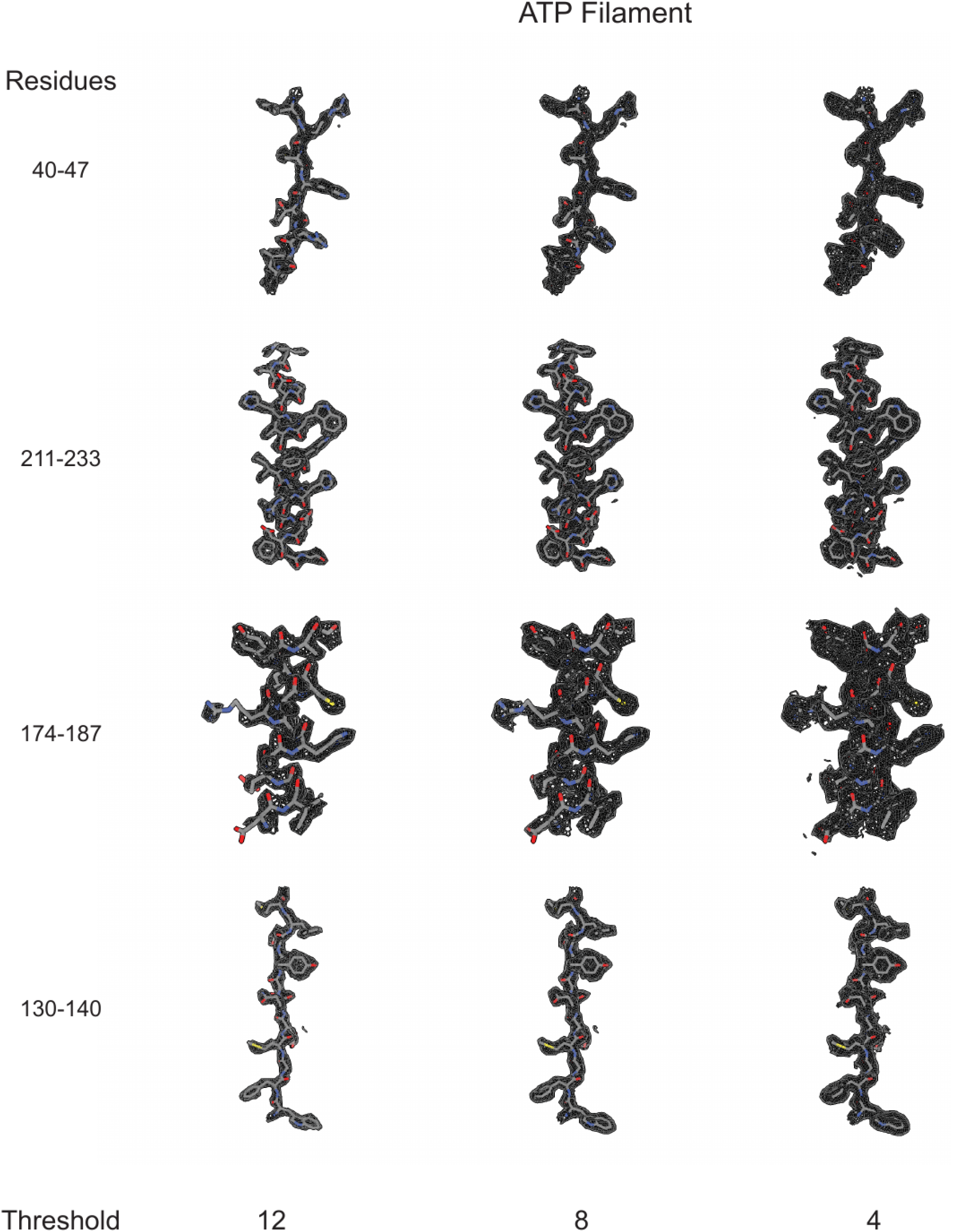
Sharpened cryo-EM density (see Methods) for the ATP-filament is presented for four regions of the map containing either β-sheet (residues 40-47 and 130-140) or α-helix (residues 174-187 and 211-233) to demonstrate side chain density and fit.

**Figure 3-figure supplement 2.**
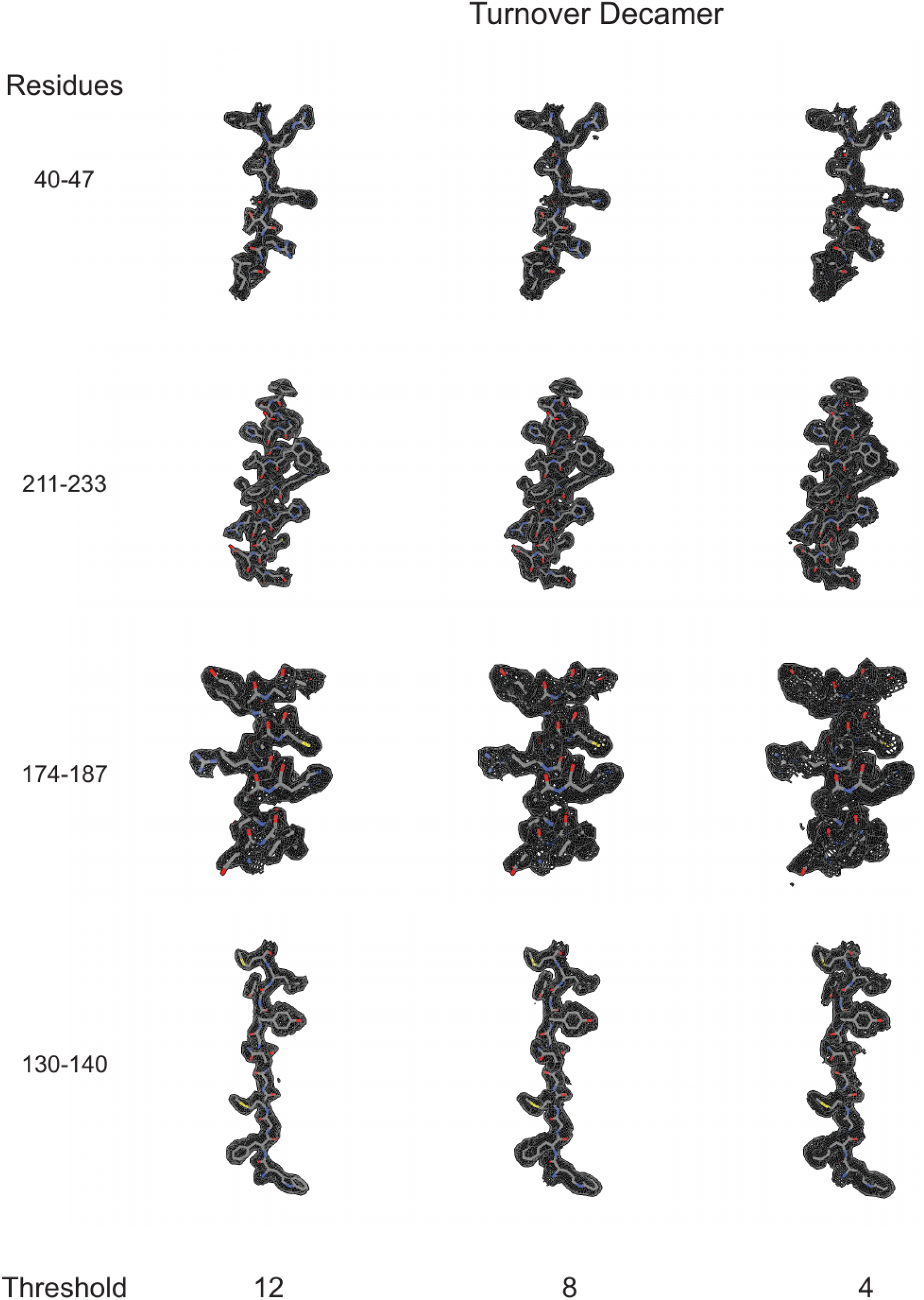
Sharpened cryo-EM density (see Methods) for the turnover decamer is presented for four regions of the map containing either β-sheet (residues 40-47 and 130-140) or α-helix (residues 174-187 and 211-233) to demonstrate side chain density and fit.

**Figure 3-figure supplement 3.**
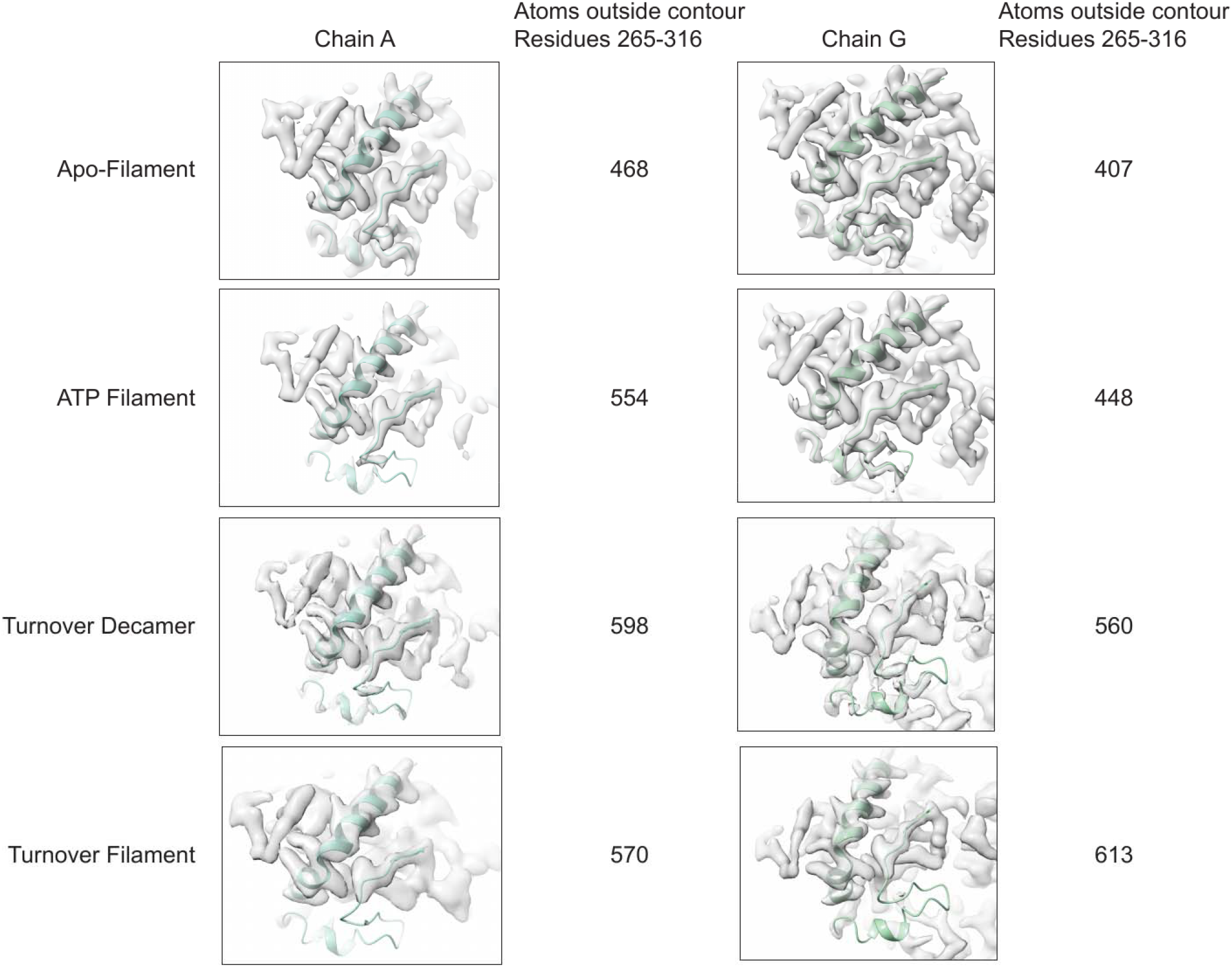
Relative loop density correlations between unsharpened cryoEM maps. Comparison of cryoEM density maps between apo-filament, ATP filament, Turnover Decamer and Turnover Filament for Chains A and G across the pentamer:pentamer interface. Density was manually thresholded in ChimeraX to be approximately similar by side change density for residues in helix 8 (265-285). A subsegment of PDB 2QC8 (residues 265-316 which includes the E305-loop (residues 289-309)) was fit into the density using the ‘Fit in Map’ tool in ChimeraX and the residues outside the contour level are reported. In general, the apo-filament map had the most density corresponding to the E305-loop, followed by the ATP filament map, Turnover Decamer map, and Turnover Filament map.

**Figure 3-figure supplement 4.**
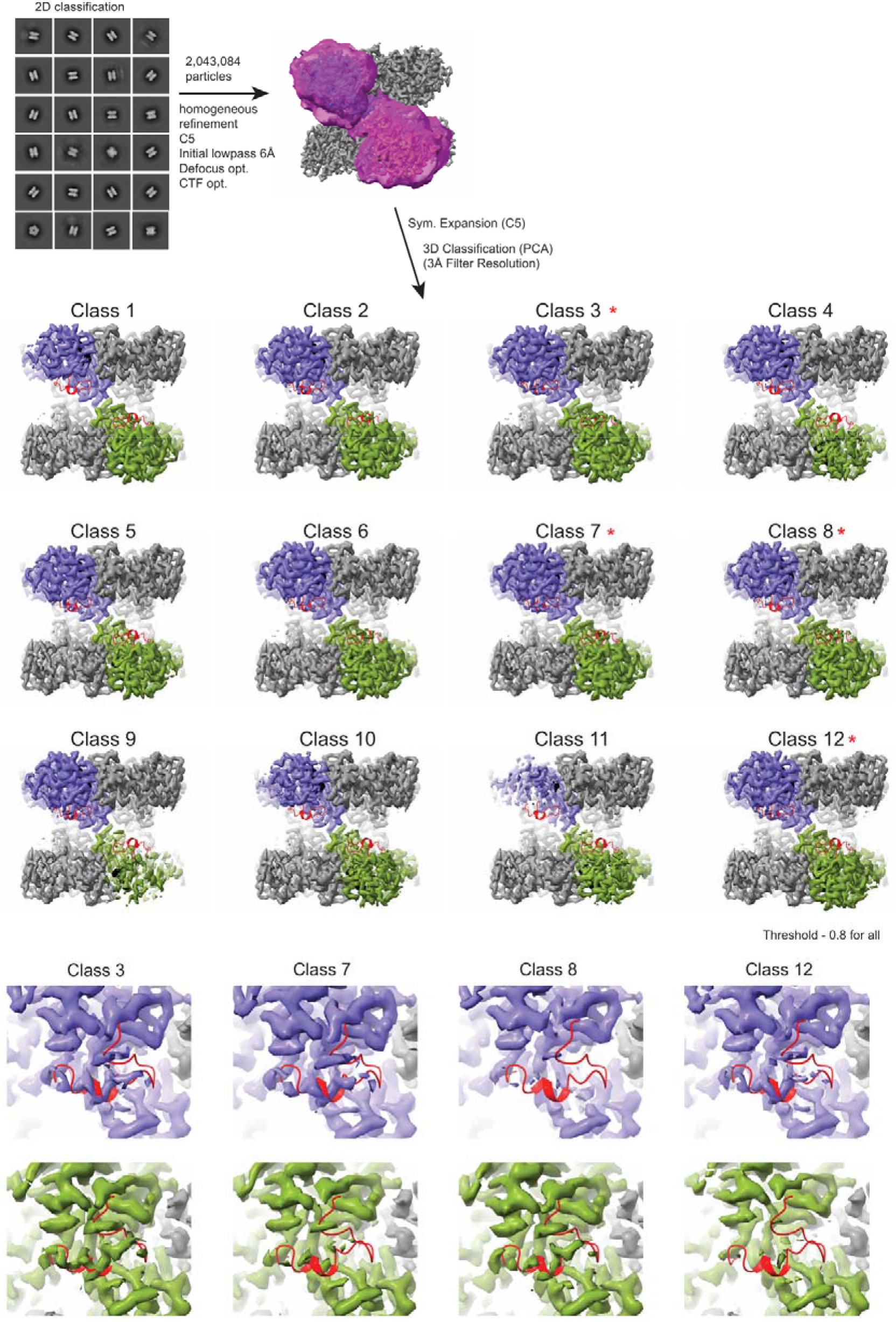
Turnover decamer focused classification reveals partial E-flap density. Pooled turnover decamer particles from the full time-resolved aggregate data were refined with C5 symmetry before symmetry expansion and focused classification on the asymmetric unit. The E-flap from PDB:2QC8 residues 290-312 (red) are fit and overlaid with each density. Four classes (3, 7, 8, and 12) were identified as having partial E-flap density, starred above and focused below, whereas all other classes were largely devoid of E-flap density.

**Figure 3-figure supplement 5.**
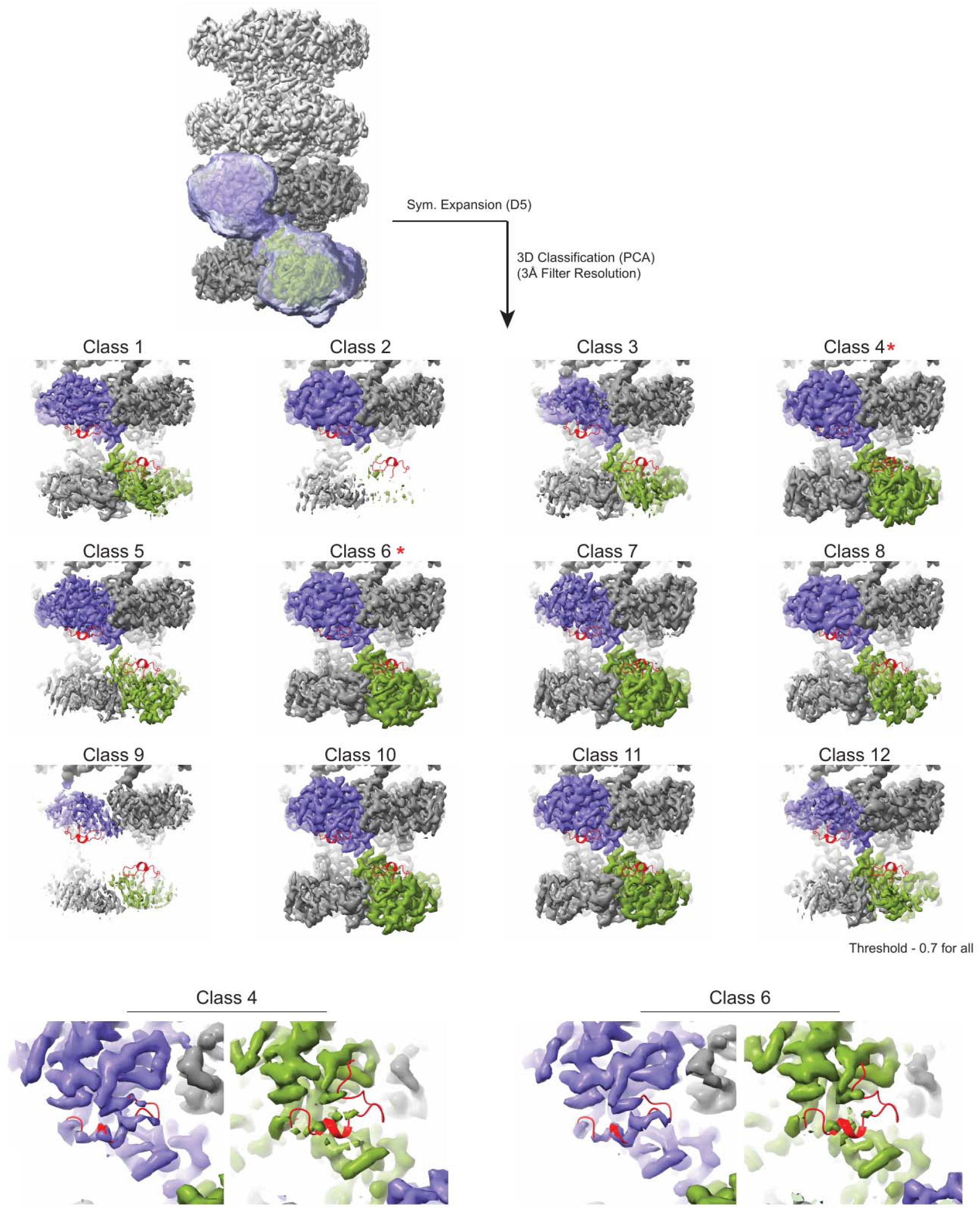
Turnover filament focused classification does not recover partial E-flap density. Turnover filament particles were symmetry expanded and subjected to focused classification on the asymmetric unit. The E-flap from PDB:2QC8 residues 290-312 (red) are fit and overlaid with each density. However, no classes were identified as having partial E-flap density whereas all other classes were largely devoid of E-flap density.

**Figure 3-figure supplement 6.**
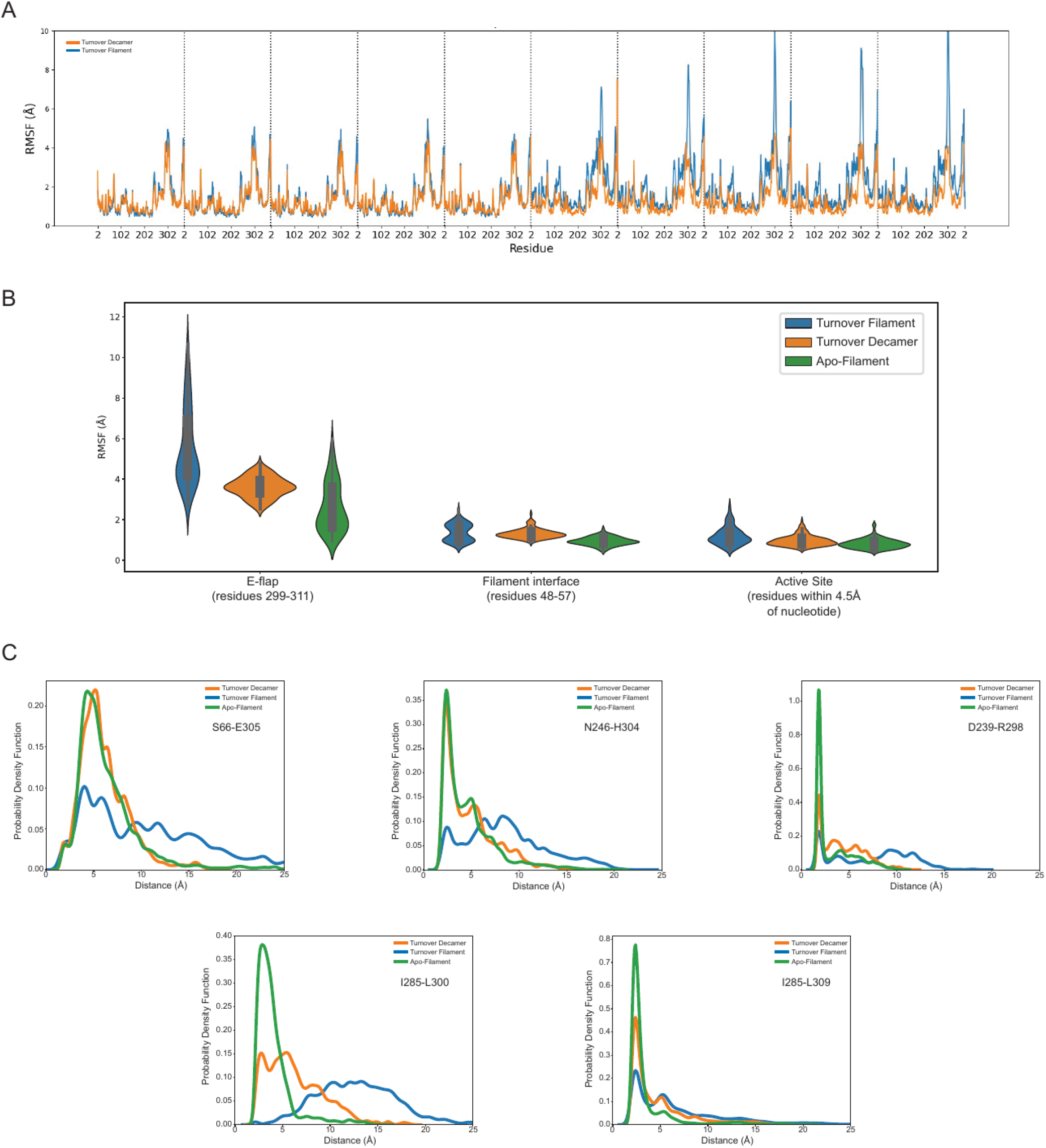
Ensemble analysis including apo-filament dataset. Top: RMSF violin plots depicting root mean square fluctuation (RMSF) of E305-loop (residues 299-311), Filament Interface loop (residues 48-57), or ‘top’ of bifunnel active site (residues within 4.5 Å of bound nucleotide) showing highest fluctuation for the E305-loop and within the E305-loop region, the highest RMSF is noted for the turnover filament followed by turnover decamer and these least RMSF for the apo-filament dataset. Both the filament interface loop and active site displayed low RMSF overall. Bottom: Similar to Figure 3 but including overlay of apo-filament ensemble data. MD-ensemble refinement distance quantification of S66-E305, N246-H304, D239-R298, I285-L300, and I285-L309. In all cases, the apo ensemble demonstrates a more stable E305-loop where all stabilizing interactions are maintained with higher probability.

**Figure 4-figure supplement 1.**
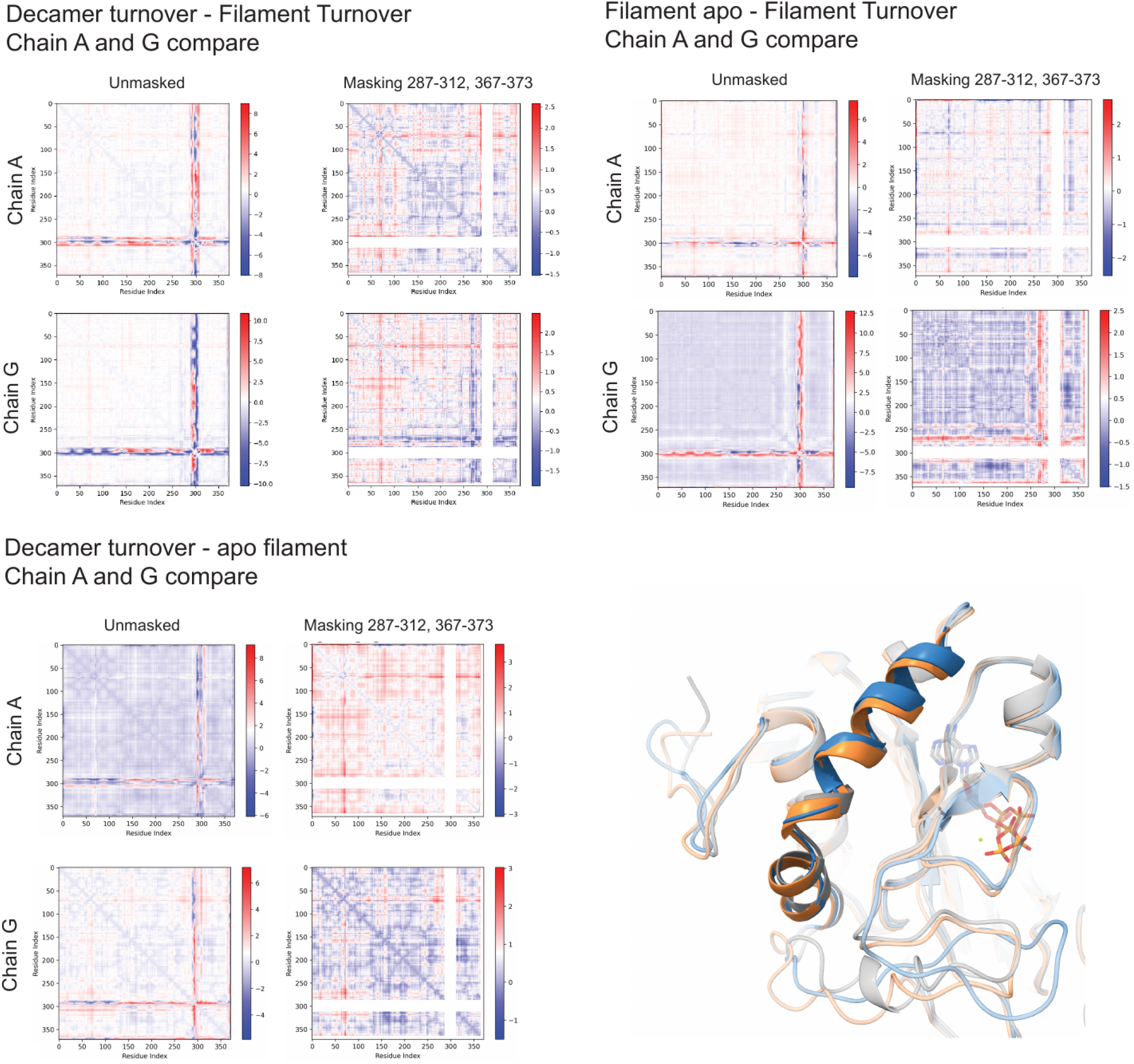
Global conformational changes most apparent in Turnover Filament structure. C-α distance difference matrix for chains A and G between Turnover Decamer and Turnover Filament models (top left), Apo Filament and Turnover-Filament models (top right), and Decamer Turnover and Apo Filament models (bottom left) depicted with and without the E305-loop (residues 289-312) and the far C-terminal peptide (369-373) omitted. Absolute distance color legend at right of each plot is in units of Å. These comparisons demonstrate that the tip of the B-grasp region (60-76) and helix 8 (residues 262-282) display conformational differences for the Filament Turnover model primarily. Bottom right: overlay of Turnover Filament (blue), Turnover Decamer (orange), and Apo Filament (gray) models highlighting helix 8 (residues 262-282) and showing a loss of helicity in part of helix 8 for Turnover Filament only.

**Figure 5-figure supplement 1.**
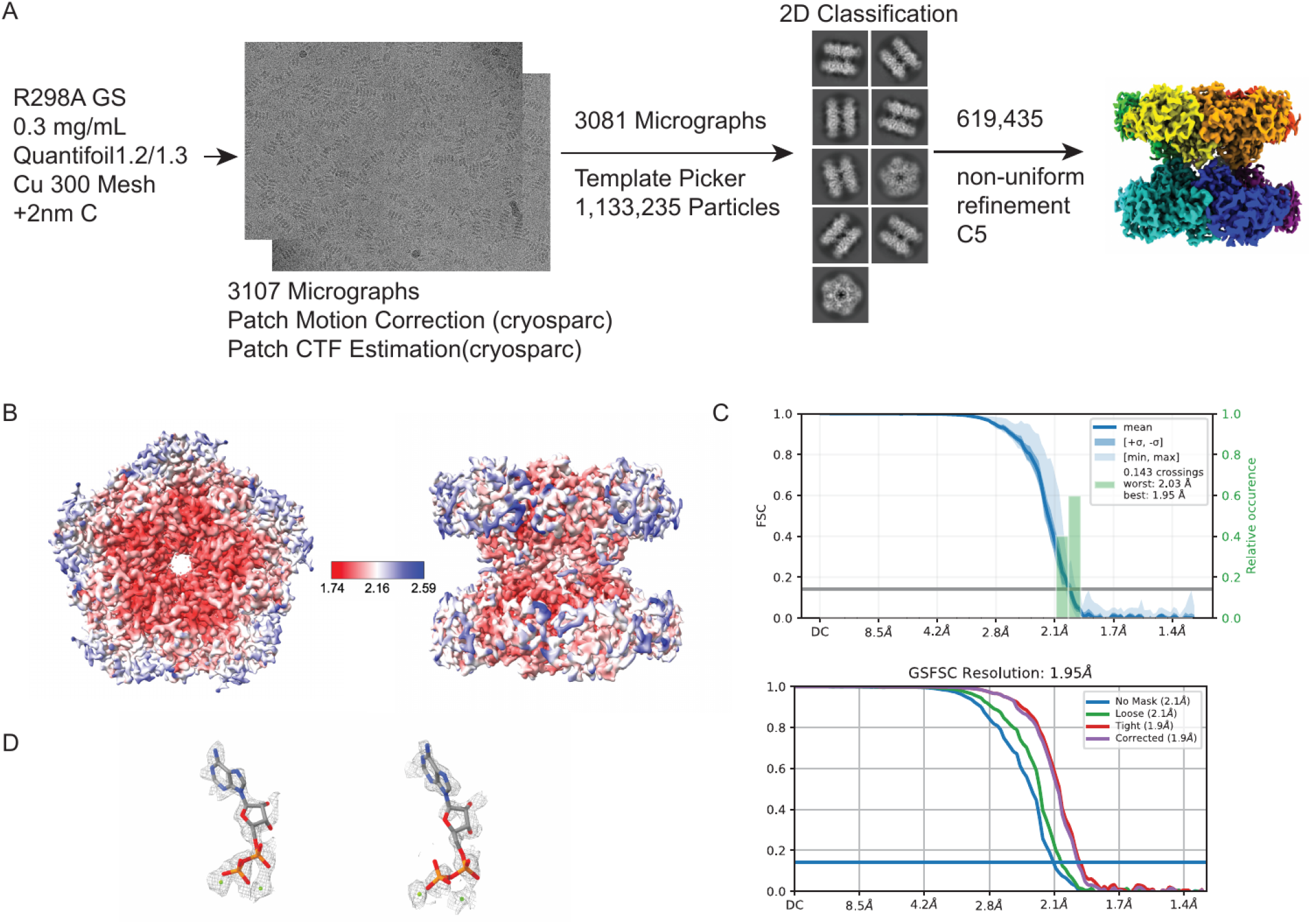
cryoEM processing pipeline for R298A turnover-decamer and comparison to wild-type turnover decamer. A) Depicted in the pipeline includes grid type, protein concentration, representative micrographs, utilized 2D classes, 3D reconstruction details, and final map (unsharpened) with associated particle counts. B) Local resolution estimation map (unsharpened). C) FSC curves for the final map. D) Active site ADP and Mg(II) density (sharpened) depiction at threshold 2.5 for chain A (left) and chain G (right).

**Figure 5-figure supplement 2.**
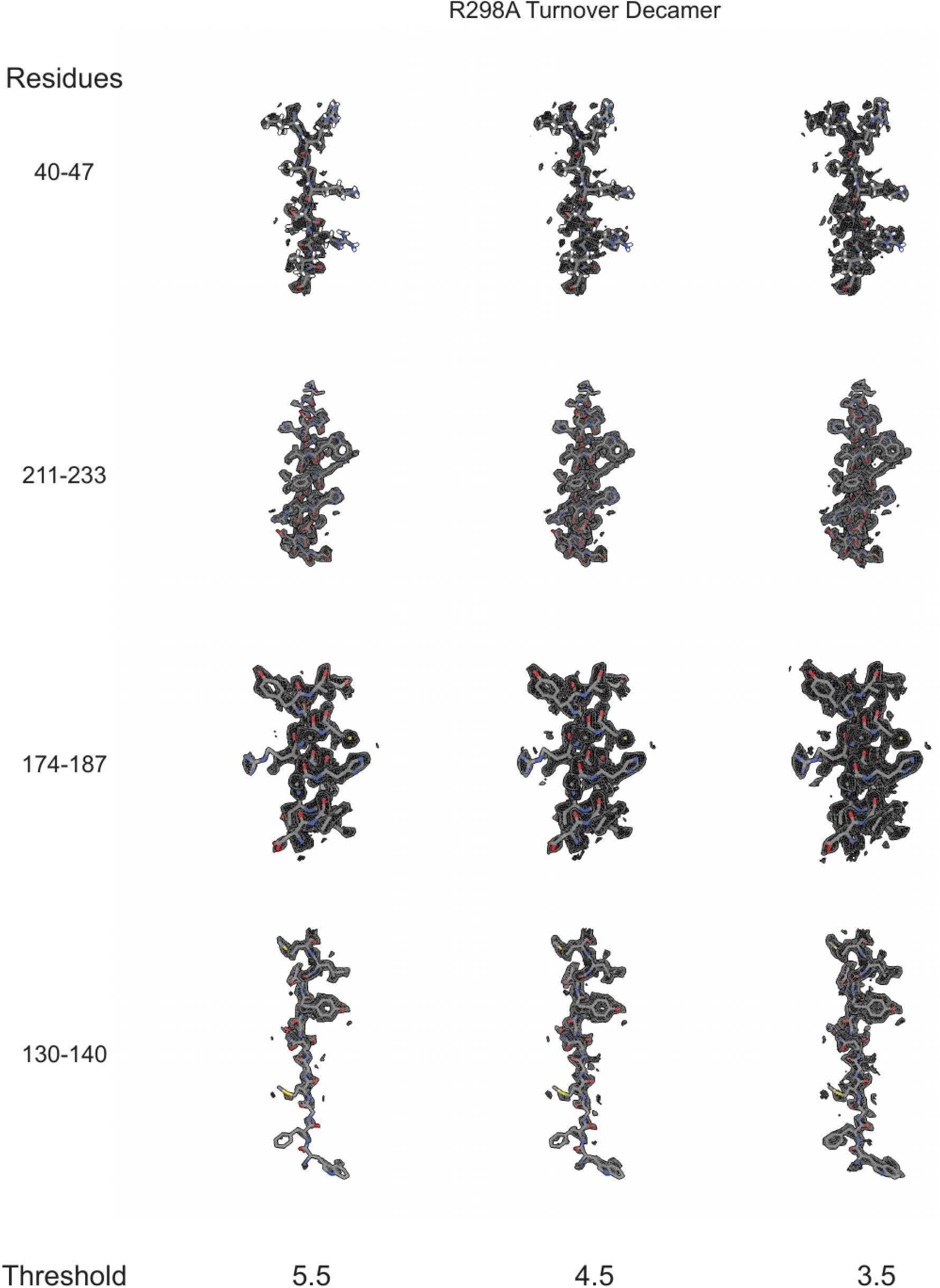
Sharpened cryo-EM density (see Methods) for the R298A turnover decamer is presented for four regions of the map containing either β-sheet (residues 40-47 and 130-140) or α-helix (residues 174-187 and 211-233) to demonstrate side chain density and fit.

**Figure 5-figure supplement 3.**
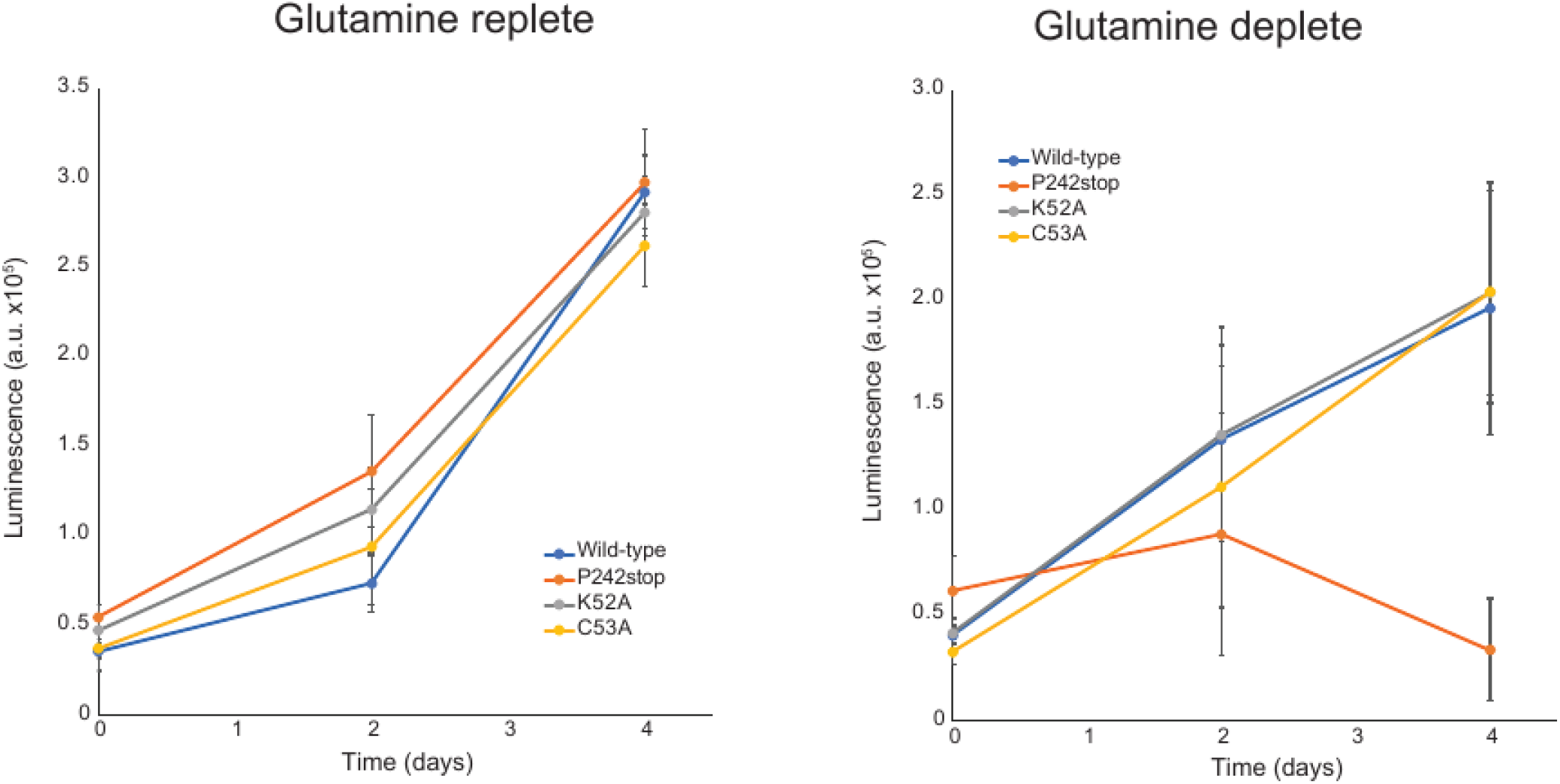
Glutamine auxotrophy determination of filament interface mutants. Luminescence derived from Promega CellTiter Glo plotted against growth time for wild-type, C53A, K52A, and P242stop variants under glutamine replete (left) or deplete (right) media conditions demonstrating similar GS-dependent growth rates for wild-type, C53A, and K52A variants, while P242stop GS could not support cell growth. Cell proliferation assays were performed in technical quadruplicate; a single clone of the landing pad cell line was generated, and a single population of each GLUL variant cell line was created for testing.

## Supplementary Material

**Supplementary Table 1.**

| Structure | Apo<br>Filament | Turnover<br>Filament | Turnover<br>Decamer | ATP<br>Filament | R298A<br>decamer |
| --- | --- | --- | --- | --- | --- |
| Ligands | Mg(II) | ADP,<br>Mg(II) | ADP,<br>Mg(II) | ATP,<br>Mg(II) | ADP,<br>Mg(II) |
| PDB code | 9OTQ | 9OTM | 9OTO | 9OTN | 9OTP |
| EMDB<br>code | EMD-70845 | EMD-70841 | EMD-70843 | EMD-70842 | EMD-70844 |
| Magnifica<br>tion | 105,000 | 106,000 | 106,000 | 105,000 | 134,000 |
| Voltage<br>(kV) | 300 | 300 | 300 | 300 | 300 |
| Electron<br>fluence<br>(e<sup>-</sup>/Å<sup>2</sup>) | 60 | 45 | 40 | 64.2 | 40 |
| Defocus<br>range<br>(µm) | 0.8-1.8 | 0.6-2.0 | 0.6-2.0 | 0.8-1.8 | 0.6-2.0 |
| Pixel size<br>(data<br>collection)<br>(Å) | 0.4215 | 0.82 | 0.86 | 0.4215 | 0.664 |
| Pixel size<br>(reconstru<br>ction) (Å) | 0.843 | 0.82 | 0.86 | 0.843 | 0.664 |
| Symmetry<br>imposed | D5 | D5 | C1 | D5 | C5 |
| Particle<br>images<br>(no.) | 112,643 | 468,883 | 2,043,084 | 310,500 | 619,435 |
| **Resolution**<br>**(0.143**<br>**FSC) (Å)** | 2.27 | 2.19 | 2.03 | 2.11 | 1.95 |
| **R.m.s.**<br>**deviation**<br>**bond**<br>**lengths**<br>**(Å)** | 0.004 | 0.004 | 0.004 | 0.004 | 0.04 |
| **R.m.s.**<br>**deviation**<br>**bond**<br>**angles (°)** | 0.942 | 0.925 | 1.018 | 0.933 | 1.026 |
| **MolProbit**<br>**y score** | 1.02 | 1.13 | 1.42 | 0.96 | 1.26 |
| **Clashscore** | 2.36 | 3.42 | 2.95 | 1.92 | 3.11 |
| **C-beta**<br>**outliers**<br>**(%)** | 0.00 | 0.00 | 0.00 | 0.00 | 0.00 |
| **Rotamer**<br>**outliers**<br>**(%)** | 0.14 | 0.14 | 1.15 | 0.32 | 0.55 |
| **Ramachandran**<br>**favored**<br>**(%)** | 98 | 98 | 96 | 98 | 97 |
| **Ramachandran**<br>**allowed**<br>**(%)** | 2 | 2 | 4 | 2 | 3 |
| **Ramachandran**<br>**outliers**<br>**(%)** | 0 | 0 | 0 | 0 | 0 |

**Supplementary Table 2.**

| Structure | Ligand | CC<br>Average ± SD (Max,<br>Min) | Qscore<br>Average ± SD (Max,<br>Min) | Bfactor<br>Average ± SD (Max,<br>Min) |
| --- | --- | --- | --- | --- |
| Apo Filament | Mg2+ | 0.52 ± 0.06 (0.60, 0.42) | 0.80 ± 0.02 (0.83, 0.77) | 94 ± 3 (99, 89) |
| ATP Filament | Mg2+ | 0.77 ± 0.07 (0.86, 0.53) | 0.88 ± 0.03 (0.93, 0.81) | 24 ± 7 (36, 10) |
| ATP Filament | ATP | 0.76 ± 0.03 (0.81, 0.71) | 0.80 ± 0.03 (0.84, 0.73) | 27 ± 9 (37, 18) |
| Turnover<br>Decamer | ADP | 0.83 ± 0.08 (0.87, 0.78) | 0.80 ± 0.01 (0.81, 0.78) | 86 ± 4 (92, 76) |
| Turnover<br>Decamer | Mg2+ | 0.71 ± 0.04 (0.79, 0.65) | 0.86 ± 0.05 (0.93, 0.75) | 74 ± 6 (85, 58) |
| Turnover<br>Filament | ATP | 0.82 ± 0.02 (0.85, 0.79) | 0.69 ± 0.03 (0.71, 0.64) | 123 ± 16 (140, 105) |
| Turnover<br>Filament | Mg2+ | 0.77 ± 0.04 (0.88, 0.60) | 0.83 ± 0.06 (0.91, 0.60) | 114 ± 22 (149, 74) |
| Turnover<br>Filament | Gln | 0.71 ± 0.04 (0.75, 0.66) | 0.51 ± 0.08 (0.59, 0.40) | 96 ± 3 (100, 93) |
| Turnover R298A | Mg2+ | 0.67 ± 0.08 (0.76, 0.51) | 0.87 ± 0.03 (0.91, 0.79) | 71 ± 9 (89, 59) |
| Turnover R298A | ADP | 0.77 ± 0.01 (0.78, 0.76) | 0.71 ± 0.03 (0.76, 0.68) | 86 ± 2 (88, 84) |

Ligand model statistics for deposited structures: Apo Filament (PDB: ), ATP Filament (PDB: ), Turnover Decamer (PDB: ), Turnover Filament (PDB: ), and Turnover R298A (PDB: ). Cross- correlation (CC) and Qscore were measured with models fit into sharpened maps in ChimeraX and Bfactor was derived from Phenix and input into deposited structures.

